# Inferring transcriptional bursting kinetics from single-cell snapshot data using a generalized telegraph model

**DOI:** 10.1101/2022.07.17.500373

**Authors:** Songhao Luo, Zhenquan Zhang, Zihao Wang, Xiyan Yang, Xiaoxuan Chen, Tianshou Zhou, Jiajun Zhang

## Abstract

**Motivation:** Gene expression has inherent stochasticity resulting from transcription’s burst manners. Single-cell snapshot data can be exploited to rigorously infer transcriptional burst kinetics, using mathematical models as blueprints. The classical telegraph model (CTM) has been widely used to explain transcriptional bursting with Markovian assumptions (i.e., exponentially distributed dwell time in ON and OFF states). However, growing evidence suggests that the gene-state dwell times are nonexponential, as gene-state switching is a multi-step process in organisms. Therefore, interpretable non-Markovian mathematical models and efficient statistical inference methods are urgently required in investigating transcriptional burst kinetics.

**Results:** We develop an interpretable and tractable model, the generalized telegraph model (GTM), to carve transcriptional bursting that allows arbitrary dwell-time distributions, rather than exponential distributions, to be incorporated into the ON and OFF switching process. Based on the GTM, we propose an inference method for transcriptional bursting kinetics using an approximate Bayesian computation framework (BayesGTM). BayesGTM demonstrates efficient and scalable estimation of burst frequency and burst size on synthetic data. Further, the application of BayesGTM to genome-wide data from mouse embryonic fibroblasts reveals that CTM would overestimate burst frequency and underestimate burst size. In conclusion, the GTM and the corresponding BayesGTM are effective tools to infer dynamic transcriptional bursting from static single-cell snapshot data.

## 1. Introduction

Gene expression is a complex biochemical reaction process with inherent stochasticity, leading to cell-to-cell variability in mRNA abundance (Elowitz, et al., 2002; Raj and Van Oudenaarden, 2008). Many experimental studies have shown that, in both prokaryotic and eukaryotic cells, the expression of most genes exhibits a stochastic burst pattern over time, characterized by silent intervals interspersed between transcriptional events of genes (Raj, et al., 2006; Yu, et al., 2006). Their burst kinetics, described by burst frequency and burst size (Rodriguez and Larson, 2020; Tunnacliffe and Chubb, 2020), are closely related to the whole molecular processes of transcriptional regulation, but the mechanism is not clear. One crucial question is how to learn and infer interpretive biological mechanisms from extensive experimental data, thus bridging the disconnect between transcriptional bursts and their underlying molecular processes, which are crucial for understanding cell-fate decisions (Eldar and Elowitz, 2010; Lammers, et al., 2020).

Addressing these questions requires visualizing transcription and measuring burst kinetics directly. A growing number of single-molecule experiments have dynamically highlighted transcriptional burst events. MS2 and PP7 imaging systems allow directly detecting the in vivo time-resolved RNA fluorescence of different genes within the same cell, revealing real-time dynamic transcriptional bursts in living cells (Bertrand, et al., 1998; Chao, et al., 2008; Chen, et al., 2020; Larson, et al., 2011). Single-molecule fluorescence in situ hybridization (smFISH) (Femino, et al., 1998; Raj, et al., 2008) quantifies the steady-state distributions of RNA in thousands of fixed single cells, from which burst parameters such as burst frequency and burst size can be inferred. However, studies based on these experimental approaches are limited to a few genes, and the burst kinetics cannot be generalized to a genome-wide perspective. Recently, single-cell RNA sequencing (scRNA-seq) (Picelli, et al., 2013; Zheng, et al., 2017) has revolutionized our understanding of cell-fate decisions and made it possible to infer the dynamic behavior of each gene from static expression distributions. To fulfill the promise of these scRNA-seq technologies, it will be crucial that mathematical models and computational methods are available to unambiguously reveal general principles of transcription on a genome-wide scale (Rodriguez and Larson, 2020).

In principle, models of gene transcription should satisfy two basic requirements. First, the gene expression model should be interpretable and mechanistic. That is, the model can offer a way to understand the mechanisms behind transcriptional bursts—for example, addressing questions such as “How do the silent transcription intervals control transcriptional bursts?” or “How transcriptional bursting relates to gene regulation?” Second, the model should be tractable. Tractability means the model can be analyzed mathematically and used to infer transcriptional bursting kinetics for large datasets. The classical telegraph model (CTM) (Peccoud and Ycart, 1995), the first rigorous mathematical treatment, connects transcription burst to stochastic gene expression. In this model, the gene switches randomly between active (ON) states and inactive (OFF), with only the former permitting transcription initiation. The CTM has been applied in the genome-wide inference of burst kinetics from scRNA-seq (Jiang, et al., 2017; Kim and Marioni, 2013; Larsson, et al., 2019; Luo, et al., 2022; Ochiai, et al., 2020; Vu, et al., 2016). For example, the inferences provided transcriptome-wide evidence that promoter elements affect burst size and enhancers control burst frequencies (Larsson, et al., 2019).

Despite the widespread use of the CTM, the model’s basic assumption – gene switching between active and inactive states at constant rates – is often unrealistic in many biological systems (Suter, et al., 2011; Tunnacliffe and Chubb, 2020; Zhang and Zhou, 2014). Mathematically, the CTM is based on the Markovian assumption that all the biochemical reaction rates involved are constant, which implies that the dwell time in each state follows an exponential distribution (Van Kampen, 1992). However, most genes have complex control processes, such as chromatin opening, recruitment of transcription factors, preinitiation complex formation, transcription initiation, as well as promoter pause and release (Fuda, et al., 2009). Such processes generate nonexponential time intervals between transcription windows. In particular, gene-state switching between active and inactive states is not a single-step manner but a multi-step process (Harper, et al., 2011; Suter, et al., 2011). This multi-step process can form a molecular memory between individual biochemical events (Zhang and Zhou, 2019), confirmed by increasing time-resolved biological experimental data (Stumpf, et al., 2017; Voss and Hager, 2014). Furthermore, this molecular memory can affect transcriptional burst kinetics (Jia and Kulkarni, 2011; Pedraza and Paulsson, 2008).

Modeling, analyzing, and inferring the molecular memory in gene-state switching is challenging. One possible way is to introduce multiple intermediate states, i.e., the promoter architecture with multiple OFF and ON states (Zhang, et al., 2012; Zhang and Zhou, 2014; Zhou and Zhang, 2012). Although the inclusion of additional gene states can improve the fit between a model and experimental data (Desai, et al., 2021; Rodriguez, et al., 2019; Zoller, et al., 2015), the difficulty in determining the number of promoter states and parameters will be detrimental to the inference of the data (Fritzsch, et al., 2018; Klindziuk and Kolomeisky, 2018; Sepúlveda, et al., 2016; Tunnacliffe and Chubb, 2020). Alternatively, one can adopt a non-Markovian modeling framework by introducing two general dwell-time distributions for OFF and ON states respectively. Importantly, the general dwell-time distributions are not limited to the exponential distribution. (Daigle Jr, et al., 2015; Kumar, et al., 2015; Schwabe, et al., 2012; Stinchcombe, et al., 2012; Zhang and Zhou, 2019a; Zhang and Zhou, 2019b). This non-Markovian model has two key advantages. First, the model is built in terms of experimentally measurable quantities and interpretable parameters, other than unobserved or inconvenient measurement gene states. Second, the model has fewer parameters due to only two gene states and therefore overcomes the difficulty in determining the number of promoter states. Despite the good properties of the non-Markovian model, it remains challenging to derive analytical solutions, develop a practical inference algorithm, and particularly infer bursting kinetics from scRNA-seq data.

In this study, we develop a statistical inference framework (BayesGTM) to infer transcriptional bursting kinetics from single-cell snapshot data. We build a generalized telegraph model (GTM) that extends the traditional exponential dwell-time distributions in ON and OFF states to arbitrary distributions. We solve the model analytically and derive the arbitrarily high-order steady-state binomial moments for mRNAs. Furthermore, we develop a statistical inference method based on approximate Bayesian computation to estimate the burst kinetics of the GTM. As a result, we show that the CTM can be misleading for inferring burst kinetics from simulation data based on the GTM. After the validation of synthetic data, the results with our inference algorithm are accurate and scalable. Finally, we apply this dynamic model and inference method to scRNA-seq data from mouse embryonic fibroblasts and find that ignoring molecular memory would overestimate burst frequency and underestimate burst size on a genome-wide scale. In conclusion, our study provides a paradigm for inferring the transcriptional bursting kinetics from single-cell snapshot data.

## 2. Model

### 2.1 Model description

Transcription occurs predominantly in episodic bursts, characterized by burst frequency and burst size (Figure 1A). The CTM is the prevailing model for describing the kinetic behavior of transcriptional bursts (Figure 1B). However, the promoter-state switching involves multiple biochemical reaction processes (Blake, et al., 2006), resulting in the number of effective states of most promoters being greater than 2 (i.e., multiple ON states and OFF states) and diverse switching between states (Sepúlveda, et al., 2016; Tunnacliffe and Chubb, 2020; Zhang, et al., 2012; Zhang and Zhou, 2014) (Supplementary Figure S1). Mapping this complex promoter architecture to the ON-OFF non-Markovian model leads to the ON and OFF dwell times being no longer exponentially distributed, as reported in previous studies (Dunham, et al., 2017; Harper, et al., 2011; Rodriguez, et al., 2019; Suter, et al., 2011).

**Figure 1.**
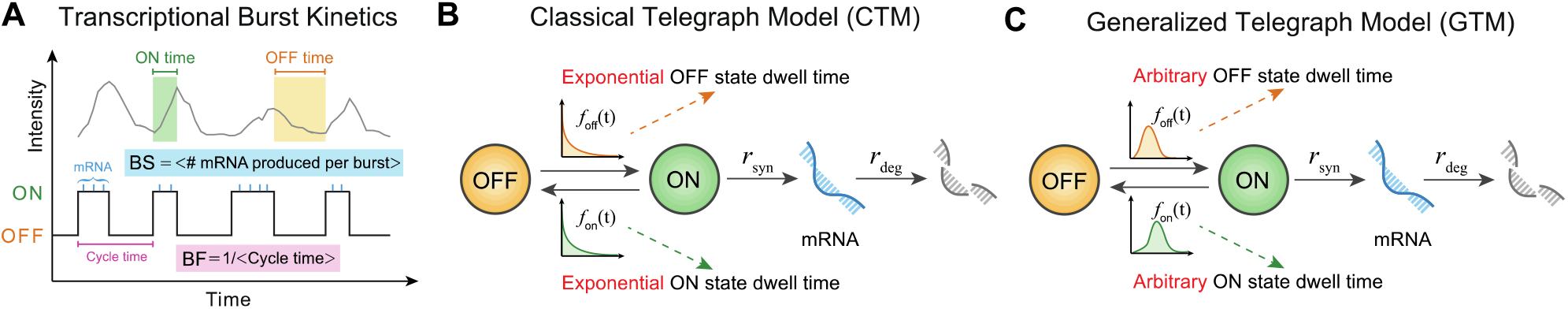
Schematic diagram burst kinetics and gene expression models. **(A)** The top panel shows a typical output of transcriptional burst in the mRNA production. The green shadow represents the time window of ON dwell time in transcription, while the orange shadow represents OFF dwell time. Correspondingly, the bottom panel represents the burst process into two states switching with each other. The short blue lines represent the transcription events during the ON state, and the purple line represents the cycle time (sum of ON and OFF time in one burst). Burst size (BS) is defined as the mean number (#) of mRNA produced per burst, and burst frequency (BF) is defined as the reciprocal of the mean cycle time. **(B)** The classical telegraph model (CTM), in which the promoter contains one OFF state and one ON state. The dwell time of the gene in these two states follows exponential distributions. The rates of mRNA synthesis and degradation are constants *r*_syn_ and *r*_deg_. **(C)** The generalized telegraph model (GTM), in which the promoter contains one OFF state and one ON state. The dwell time of the gene in these two states follows arbitrary distributions *f*_off_ (*t*) and *f*_on_ (*t*). The rates of mRNA synthesis and degradation are constants *r*_syn_ and *r*_deg_.

To make this idea precise, we consider a more general stochastic transcription model, called GTM, as illustrated in Figure 1C. We assume the dwell times in OFF and ON states, two random variables *τ*_off_ and *τ*_on_, follow arbitrary probability distributions, denoted by *f*_off_ (*t*) and *f*_on_ (*t*), rather than the limited exponential distributions. The mRNA synthesis and degradation process are assumed to be Markovian, i.e., exponential distributions of transcription waiting time *f*_syn_ (*t*) and mRNA’s lifetime *f*_deg_ (*t*). Specifically, 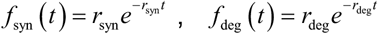, where *r*_syn_ and *r*_deg_ are the mean rate of mRNA synthesis and degradation, respectively. The reaction scheme is summarized by the reaction diagram:

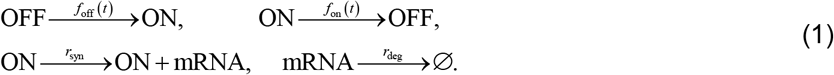

### 2.2 Burst kinetic of GTM

The burst size and burst frequency (or the burst cycle time) are the two most critical parameters to characterize burst kinetics. First, we derive the probability distributions and their statistics for the burst size and cycle time with the GTM (Figure 1A). Burst frequency describes the average number of bursts that occurred per unit time, i.e., the reciprocal of the mean cycle time. According to the definition of cycle time, which is the summation of the *τ*_off_ and *τ* _on_, the distribution of cycle time is the convolution of the distributions *f*_off_ (*t*) and *f*_on_ (*t*), i.e., (*f*_off_ * *f*_on_)(*t*). Consequently, we can obtain the expression of burst frequency for GTM (see Supplementary Note 1.1 for a detailed derivation)

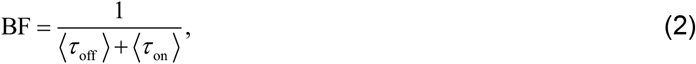

Where 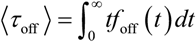 and 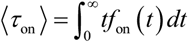 are the mean OFF and ON state dwell time, respectively.

Burst size describes the average number of mRNA molecules produced per burst. We first derive the distribution of transcription-event numbers per burst. For the exponential transcription process with rate *r*_syn_, the probability of the occurrence of *x* transcription events conditioned on a fixed duration time *t* of the ON state is a time-dependent Poisson distribution 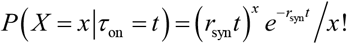. Then the probability of the transcription-event number per ON state period, denoted by *P* (*X*), can be computed by the total probability theorem 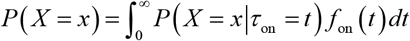. Therefore, we obtain the burst size for GTM by some algebraic calculations (see Supplementary Note 1.2 for a detailed derivation)

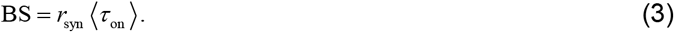

Next, we confirm the necessity of the GTM with the help of the obtained burst kinetics (Eq. (2) and Eq. (3)) of the GTM. From the principle of parsimony, we know that the burst size and frequency can also be directly computed with the CTM (Kim and Marioni, 2013; Larsson, et al., 2019; Ochiai, et al., 2020). A natural and important question is whether the CTM quantitatively enough describes the burst kinetics (burst size and burst frequency). We answer it by performing inference on simulation data from GTM. First, we repeatedly generate synthetic scRNA-seq data using the simulation algorithm of GTM (see Supplementary Table S1). In the GTM, molecular memories are characterized by Gamma distributions (Pedraza and Paulsson, 2008), i.e., 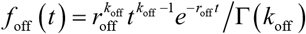 and 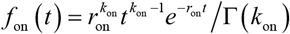 where Γ (·) is gamma function, *r*_off_ and *r*_on_ are the rate-parameters of state switching, and *k*_off_ and *k*_on_ are the number of possible reaction steps from OFF to ON and vice versa. Then, we use the CTM to estimate the burst kinetic parameters of the synthetic data via the maximum likelihood estimation approach. As a result, we find that although the CTM can well fit the gene expression distributions generated by the GTM (Figure 2A and 2E), it cannot accurately describe the dwell-time distributions (Figure 2B, C, F, G), further leading to the erroneous estimations of burst frequency and burst size (Figure 2D and 2H). These results suggest that when we use the non-time-resolved single-cell snapshots data to infer burst kinetics, different dwell-time distributions may yield identical gene expression distributions, which leads to misunderstandings of burst kinetics inferred from the CTM. Therefore, it is necessary to develop a scalable statistical inference approach to accurately estimate the burst kinetics from genome-wide scRNA-seq data based on the GTM.

**Figure 2.**
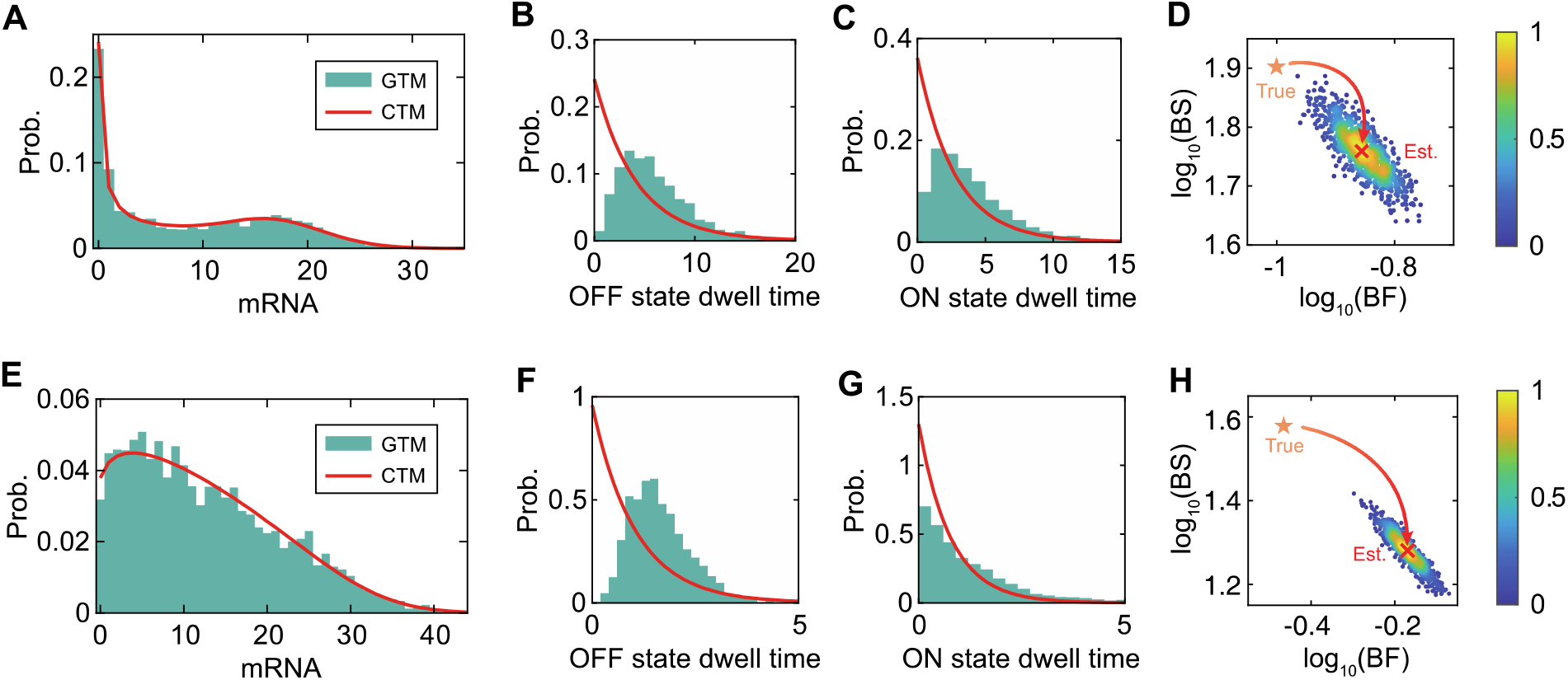
The CTM fits well the data generated by the GTM, but misleads the burst kinetics. **(A)** Histogram represents the data generated from the GTM with parameters *k*_off_ = 3, *r*_off_ = 0.5, *k*_on_ = 2, *r*_on_ = 0.5, *r*_syn_ = 20, *r*_deg_ = 1. The solid red line represents the mRNA distribution estimated from the CTM. **(B)** Histogram shows the simulated data of OFF-state dwell time from the GTM. The solid red line represents the distribution of OFF-state dwell time estimated from the CTM. **(C)** Results similar to **(B)** corresponds to the ON-state dwell time. **(D)** Scatter plots of transcriptional burst frequency (BF) and burst size (BS) were obtained from 1000 repeats of GTM simulations and CTM estimations. The red cross represents the most probable parameters obtained by using smooth kernel density for the burst parameters estimated by the CTM, while the orange star represents the true burst parameters calculated from the GTM. **(E-H)** The results of another example with the parameters *k*_off_ = 5, *r*_off_ = 3, *k*_on_ = 1, *r*_on_ = 0.8, *r*_syn_ = 30, *r*_deg_ = 1.

### 2.3 Model analysis

To perform statistical inference on the burst kinetics by a steady gene-expression distribution from static scRNA-seq data, we theoretically solve the statistical properties of the transcriptional process described in Eq. (1) using a supplementary variable method (Alfa and Rao, 2000; Cox, 1955). Denote by *M* (*t*) the number of mRNA molecules at time *t* and *G* (*t*) the gene state at time *t*. *E*_off_ (*t*) (*E*_on_ (*t*)) is defined as the elapsed time since the gene switches to the OFF (ON) state at time *t* respectively. Then, {*M* (*t*),*G*(*t*), *E*_off_ (*t*), *E*_on_ (*t*);*t* ≥ 0} is a continuous-time Markov process. Let *p*_off_ (*n,τ, t*) and *p*_on_ (*n,τ, t*) be the probability density functions (PDF) that *n* mRNA molecules are produced and the elapsed time is *τ* in OFF and ON state at time *t* respectively, and we have

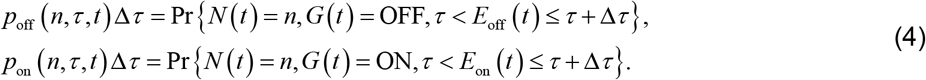

According to the relationship between the states of the system at time *t* and *t* + Δ*t*, we obtain the following Chapman–Kolmogorov backward equations,

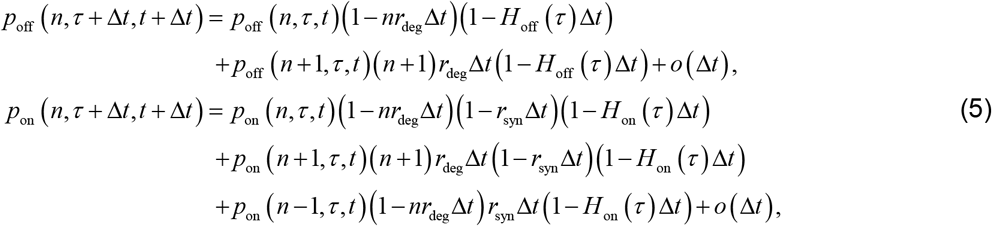

where *H*_off_ (*τ*) = *f*_off_ (*τ*) *S*_off_ (*τ*) is a hazard rate function with the survival function 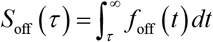.

The definitions of *H*_on_ (*τ*) and *S*_on_ (*τ*) are similar.

Next, we focus on steady-state distributions. If the stationary distribution of *p*_off_ (*n,τ, t*) and *p*_on_ (*n,τ, t*) exist (above simulation has verified this point), and are denoted by *p*_off_ (*n,τ*) and *p*_on_ (*n,τ*) respectively, Eq. (5) converts to the following stationary chemical master equation in the limit of small Δ *t* and large *t*,

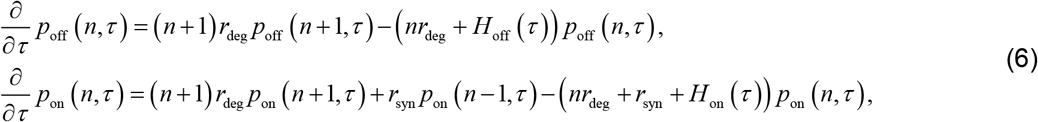

with the integral boundary conditions 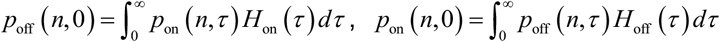.

Based on Eq. (6) with its boundary conditions, we use the binomial moment method (Zhang, et al., 2016) to calculate the mRNA stationary distribution 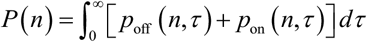 and its statistical characteristics. Binomial moments of the mRNA stationary distribution are defined as 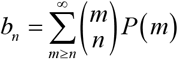, where the symbol 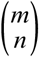 represents the combinatorial number. Binomial moments converge to zero as their orders go to infinity, and can be used to reconstruct *P* (*n*) by 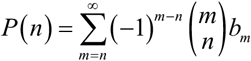. After some algebra of Eq. (6), we can obtain the *n* -th binomial moment of mRNA in a recursive form (see Supplementary Note 2 for a detailed derivation)

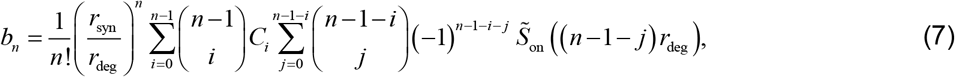

where

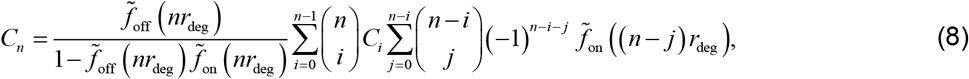

for *n* = 1, 2,…. Here *C*_0_=(⟨*τ*_off_ ⟩)+(⟨*τ*_on_ ⟩)^−1^ is equal to the burst frequency. 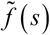 represents the Laplace transform of function *f* (*t*). Especially, 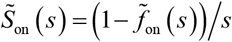 and 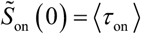. According to the relationship between binomial moments and central moments and Eq. (7) and Eq. (8), we obtain the mean and noise of mRNA expression

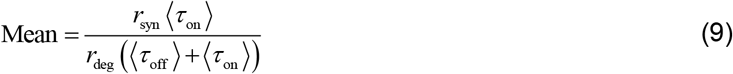

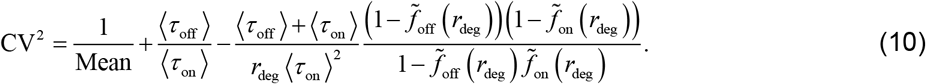

Note that if let *τ* _off_ *r*_deg_ and *τ*_on_ *r*_deg_ be the rescaled random variables for OFF-state and ON-state dwell times, and *r*_syn_ *r*_deg_ be the rescaled mean synthesis rate, the mean expression not only equals the product of the mean synthesis rate and the stationary probability of ON state, but also the product of burst size and burst frequency.

Finally, we consider two specific cases of the GTM. First, if the ON-state dwell time is exponential, i.e., 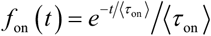, the GTM and corresponding results reduce to the previous conclusions (Shi, et al., 2020) (see Supplementary Note 3.1). Further, if the OFF-state dwell time is also exponential, i.e., 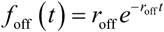, the GTM reduces to the CTM. In this model, the mean activation and inactivation rates are *r*_off_ = ⟨*τ*_off_⟩^−1^ and *r*_on_ = ⟨*τ*_on_⟩^−1^, respectively. Notably, the expression for the *n* -th binomial moment of mRNA can be simplified as follow (see Supplementary Note 3.2),

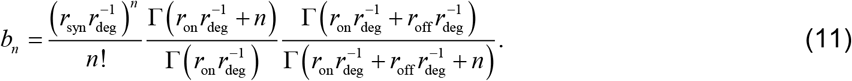

Using the reconstruct formula, we obtain the mRNA stationary distribution for CTM (see Supplementary Note 3.2), consistent with previous results (Peccoud and Ycart, 1995; Raj, et al., 2006).

## 3. Binomial moment-based inference

In this section, we will develop a statistical inference method based on binomial moments for GTM. First, we assume that the number of cells in the experiment is large enough to allow us to acquire a steady expression distribution of a specific gene. This assumption is reasonable in scRNA-seq data and has been widely applied to genome-wide studies (Larsson, et al., 2019; Luo, et al., 2022). Next, with the steady distribution of each gene from scRNA-seq data as a bridge, we develop a computation framework (BayesGTM) that uses an approximate Bayesian computation (ABC) approach (Franks, 2020) to infer reliable parameter sets for our GTM (see Supplementary Table S2).

### 3.1 BayesGTM framework

BayesGTM is a statistical framework that combines the ABC approach to estimate the Bayesian posterior probability *π* (**θ**❘ **y**_obs_) of the model parameter-vector (**θ**) given the observed scRNA-seq data (**y**_obs_). The prior information related to **θ** is denoted as the prior distribution *π* (**θ**), which will be iteratively updated through computing the likelihood function *p*(**y**_obs_ ❘**θ**) of the GTM. Using Bayes’ theorem, the resulting posterior distribution is given by

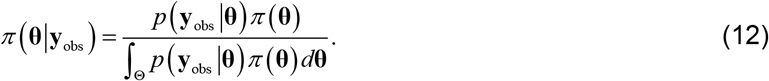

Ideally, we can perform the inference methods relying on iterative likelihood function maximization. However, the approximation of the mRNA distribution of the GTM is computationally prohibitive, making it impossible to evaluate the likelihood function directly.

Alternatively, we resort to a “likelihood-free” Bayesian approach by computing only low-order moments instead of the whole probability distribution according to the obtained binomial moments (Eq. (7)). Specifically, we use the sequential Monte Carlo ABC (Sisson, et al., 2007) (see Supplementary Table S2), a variant of ABC, to implement the statistical inference of our model. ABC allows to accept the parameters that make the simulated data and the observed data sufficiently close in distribution and to estimate the posterior distribution *π* (**θ❘y**_obs_) of the parameters through numerous simulations. First, given a dynamic model (Figure 3A), we sample a candidate parameter vector **θ** from the prior distribution *π* (**θ**) (Figure 3B), and simulate a dataset **y**_model_ from the GTM. Then, we check whether the distribution of simulated data approximates the observed data **y**_obs_ (Figure 3D) by predefining three extra parameters: (1) Summary statistics **s** (**y**) for sufficiently representing data; (2) Discrepancy metrics *ρ* (·,·) for measuring the distance between summary statistics of observed data **s**_obs_ and of simulated data **s**_model_ from the GTM (Figure 3E); (3) Threshold *ε* for controlling acceptable errors (Figure 3F). Notice that the low threshold *ε* of ABC promises a good approximation of *π* (**θ❘ y**_obs_), but also imposes a huge computational cost and low-rate acceptances. To avoid the difficulties of ABC in terms of computational power and convergence, we use the sequential Monte Carlo ABC.

**Figure 3.**
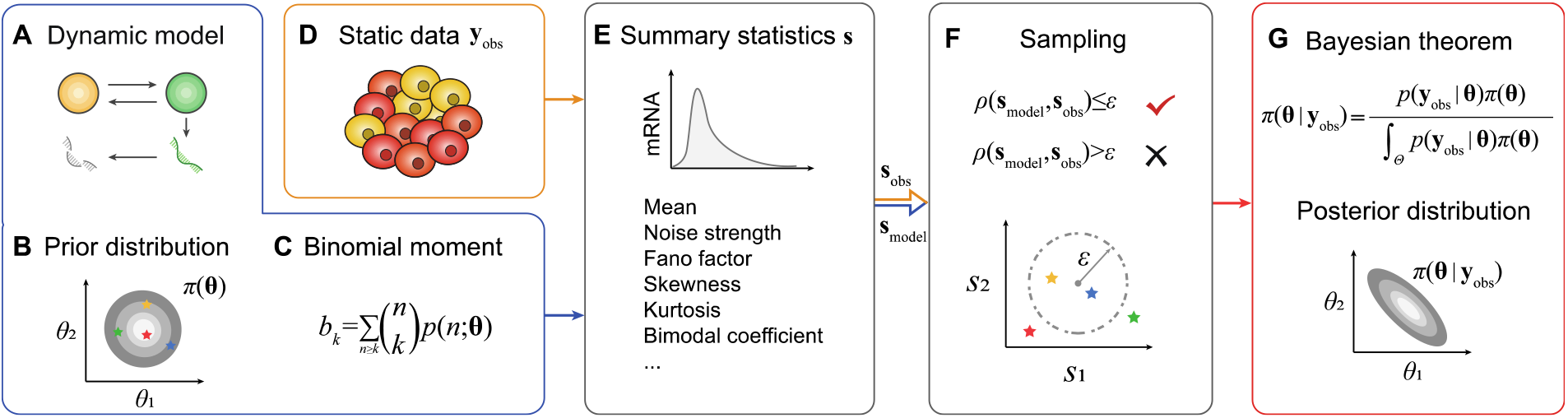
BayesGTM inference procedure. Given a dynamic model **(A)**, the parameter **θ** is sampled from the prior distribution *π* (**θ**) **(B)**, and then the theoretical binomial moment **(C)** is calculated to compute the summary statistic **s**_model_ **(E)**. The static single-cell snapshot data **y**_obs_ **(D)** can be used to calculate the summary statistic **s**_obs_ **(E)**. The sampled **θ** is accepted by comparing whether *ρ* (**s**_obs_, **s**_model_) is less than the threshold *ε* **(F)**. As the output of BayesGTM, the posterior distribution *π* (**θ❘ y**_obs_) of the parameters is obtained by Bayes’ theorem **(G)**.

Sequential Monte Carlo ABC adds a sequence of threshold values {*ε*_0_, *ε*_1_,…*ε*_*T*_} satisfied the condition *ε*_0_>…> *ε*_*T*_ ≥ 0 and thus constructs a sequence of the intermediate posterior distributions { *f*_*t*_ (**θ❘** *ρ* (**s**_obs_, **s**_model_) ≤ *ε*_*t*_)}, *t* ϵ{0,1,…,*T*}, where *f*_0_ (**θ**) = *π* (**θ**). When the (*t* −1) -th iteration is over, we can obtain the *f*_*t* −1_ (**θ❘** *ρ* (**s**_obs_, **s**_model_) ≤ *ε*_*t*−1_). In the *t* -th iteration, we first sample the parameters **θ**_*t* −1_ randomly from *f* _*t* −1_(**θ❘** *ρ* (**s**_obs_, **s**_model_) ≤ *ε* _*t* −1_) and sample a **θ**^*^ from the proposal distribution *K* _*t*_ (**θ❘ θ** _*t* −1_). Then we accept the **θ**^*^ if *ρ* (**s**_obs_, **s**_model_(**θ**^*^)) ≤ *ε* _*t*_. After repeating the above procedures in this iteration, we get the posterior distribution *f*_*t*_ (**θ❘** *ρ* (**s**_obs_, **s**_model_) ≤ *ε*_*t*_). In theory, *f*_*T*_ (**θ❘** *ρ* (**s**_obs_, **s**_model_) ≤ *ε*_*T*_) = *π* (**θ❘ y**_obs_) when *ε*_*T*_ → 0. Finally, we can obtain the posterior distribution *π* (**θ❘ y**_obs_) (Figure 3G).

### 3.2 BayesGTM inference procedure

#### Summary statistics

BayesGTM inference requires a summary statistic **s** (**y**) to reduce the high-dimensional data to low-dimension features to compare whether the distribution of simulated data **y**_model_ from the GTM is close to the observed data **y**_obs_. Here, we choose six commonly used moment statistics for inference (Fröhlich, et al., 2016; Zechner, et al., 2012) that are important to characterize the shape of gene expression distribution as the summary statistics: (1) The mean value *μ*_1_ is the most commonly used indicator in statistics, and it represents the average level of mRNA expression in scRNA-seq data. (2) The noise strength is a measurement of the dispersion of the probability distribution, defined as 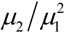 where *μ*_2_ is the variance. (3) The Fano factor is another statistic that measures the dispersion of a probability distribution relative to a Poisson distribution, defined as *μ*_2_ *μ*_1_. (4) The skewness is a description of the symmetry of the distribution. Skewness is defined as 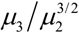, where *μ*_3_ is the third central moment. (5) The kurtosis describes whether the peak of the distribution is abrupt or flat, which is defined as 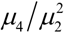, where *μ*_4_ is the fourth central moment. (6) The bimodality coefficient can describe the bimodal distribution (Ellison, 1987), which is usually a critical feature in a dynamical system. The values of the bimodality coefficient range from 0 to 1, and values greater than 5/9 may indicate a bimodal or multimodal distribution.

Crucially, BayesGTM used the binomial moment approach theoretically provides an efficient method to directly compute the theoretical summary statistic for a given parameter **θ** of the GTM. Precisely, we can calculate the central moments with the binomial moment:

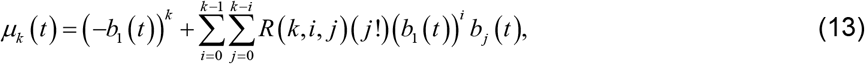

in which 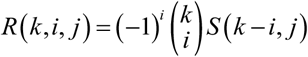 with 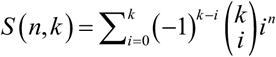 being the Stirling number of the second kind. Therefore, the above summary statistics can be expressed by binomial moments as follows

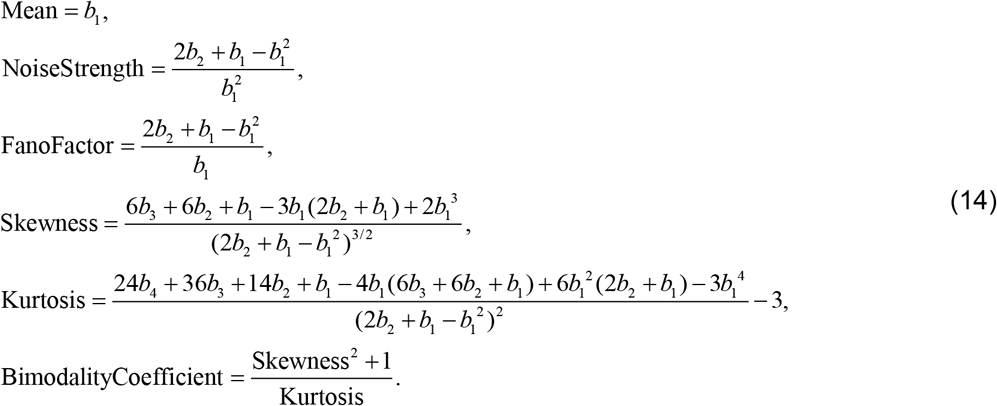

It should be noted that we can extend the summary statistics to higher-order moments because our binomial moments can compute arbitrary high-order moment statistics.

To assess the sensitivity of the summary statistics, we investigate the influence of the model parameters (*k*_on_, *k*_off_, *r*_on_, and *r*_off_) on the six summary statistics through the simulation algorithm of GTM (Supplementary Table S1) and the theoretical results (Eq. (14)). The results show the six summary statistics are sensitive to the parameters of dwell times (Supplementary Figure S2), implying the rationality of statistics selection.

#### Discrepancy metrics

To eliminate the influence of absolute size between different summary statistics, we take the natural logarithm of the data instead of computing the Euclidean distance (Desai, et al., 2021; Lenive, et al., 2016)

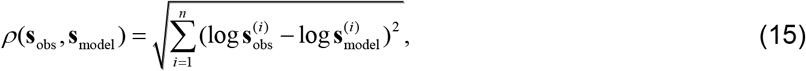

where **s**^(*i*)^ is the *i* -th component of the summary statistics vector. Note that the logarithm transformations of data do not change the data properties and correlation, but the scale is compressed. In addition, these transformations can make the data more stable and weaken the effects of collinearity and heteroscedasticity on the model.

#### Acceptance threshold

Generally, the acceptance thresholds {*ε*_0_, *ε*_1_,…*ε*_*T*_} are usually determined by experience. However, the algorithm causes a waste of computational resources in this iteration if the default threshold is greater than the maximum distance obtained by sampling in the last iteration. Therefore, we adopt a strategy of adaptive acceptance thresholds to prevent this situation. For the first iteration, we use a large threshold *ε*_0_ to quickly accept 10000 coarse parameter samples, and then select the first 1000 parameter samples with the smallest discrepancy between **s**_obs_ and **s**_model_ as input for the next iteration. For the *t* –th iteration, the acceptance threshold *ε*_*t*_ is set to the median of the discrepancies of the results obtained from the (*t* −1) -th iteration.

#### Prior distribution

As required in Bayesian inference, we should set prior distribution *π* (**θ**), allowing for the initial parameter sampling **θ** in the BayesGTM inference procedure. In the GTM, we assume that the mRNA decay rate *r*_deg_ = 1 and the dwell-times in the ON state and OFF state are Gamma distributed, i.e., 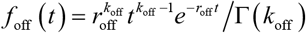 and 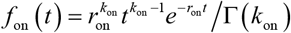, respectively. The parameter-vector is **θ =** [*k*_off_, *r*_off_, *k*_on_, *r*_on_, *r*_syn_]. We set the prior distributions of *k*_off_ and *k*_on_ follow the uniform distribution *U* [0, 5] and the prior distributions of *r*_off_ and *r*_on_ follow the log-uniform distribution from interval [−1,1] based on 10. In addition, the prior distribution of transcriptional rates *r*_syn_ is *U* [0, 50].

#### Proposal distribution

In the BayesGTM inference procedure, the parameter **θ**^*^ sampled in *t* -th iteration is based on small perturbations around the parameter **θ**_*t* −1_ sampled from the *f*_*t* −1_ (**θ❘** *ρ* (**s**_obs_, **s**_model_) ≤ *ε*_*t*−1_). Therefore, we use lognormal distribution as the proposed distribution *LN* (**θ**;**θ**_*t* −1_, *σ*) for sampling all parameters in the *t* –th iteration.

## 4. Results

### 4.1 Synthetic data

We first use synthetic scRNA-seq data from the GTM to verify whether the BayesGTM can accurately estimate the burst kinetics of the gene expression. We apply the simulation algorithm for the GTM with given parameters (see Supplementary Table S1) to generate mRNA distribution. Figure 4A shows the GTM that the dwell times for OFF and ON states are nonexponential distributions and have a bimodal distribution at a steady state, which has important significance in biological systems (Ochab-Marcinek and Tabaka, 2010) and is a common observation in scRNA-seq data (Sarkar and Stephens, 2021). Having obtained 1000 samples by the BayesGTM inference procedure, we find that the posterior distribution of transcriptional rate *r*_syn_ can accurately estimate the true parameter value (Figure 4B). Importantly, we show that the dwell time’s first-order moment information (mean) can be estimated accurately (Figure 4C). And the burst frequency and burst size can also be estimated accurately (Figure 4D) (although the individual parameters are unidentifiable in large part of parameter space). In another example, the distribution of GTM is unimodal, and the burst kinetics can also be accurately inferred (Figure 4F-H).

**Figure 4.**
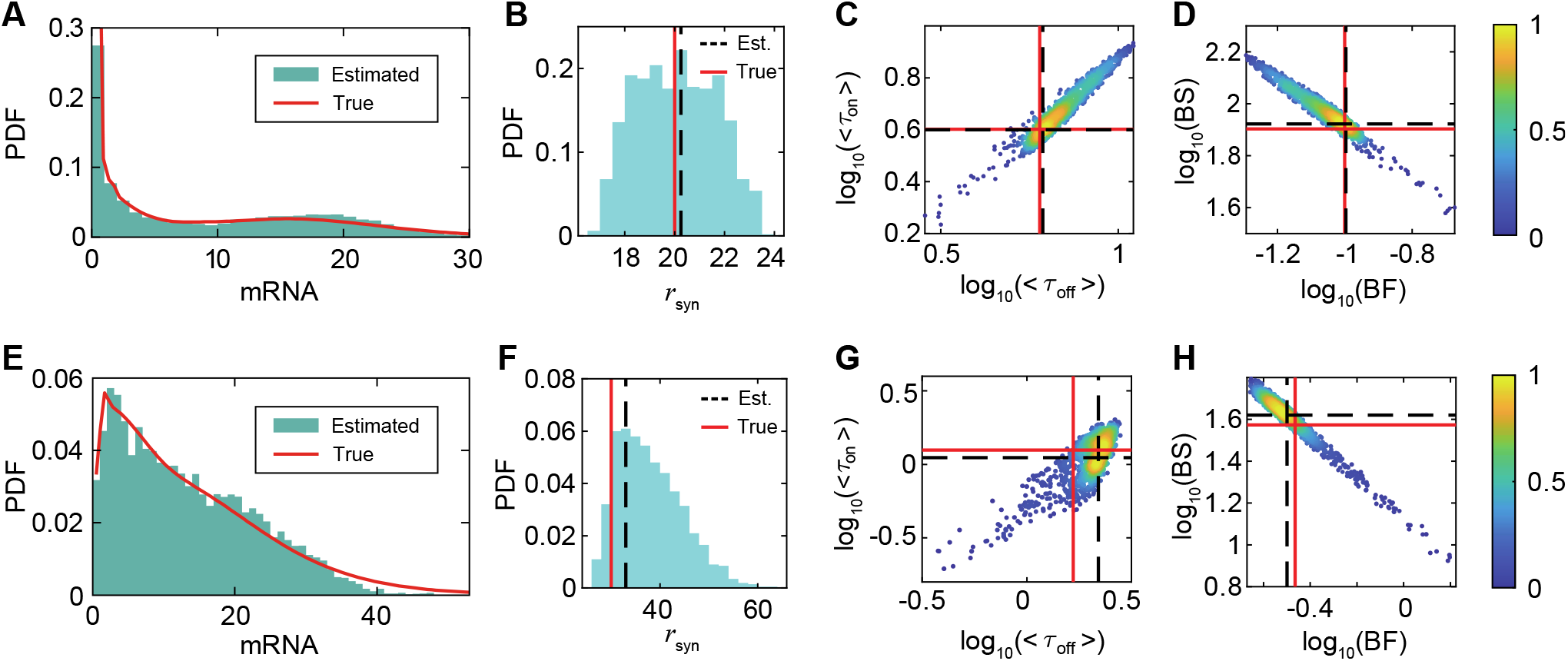
Validation of inferring burst kinetics on synthetic scRNA-seq data. **(A)** Comparison between synthetic scRNA-seq data for inference with parameters *k*_off_ = 3, *r*_off_ = 0.5, *k*_on_ = 2, *r*_on_ = 0.5, *r*_syn_ = 20, *r*_deg_ = 1 (red solid line, the PDF fitted by kernel smoothing function) and simulation data of inferred parameters (green histogram). **(B)** Marginal posterior distributions (blue histogram) of transcriptional rate *r*_syn_ estimated by BayesGTM, where the red solid line corresponds to the true parameter and the black dash line to the mode of estimated parameter. **(C)** The scatter plot shows the two-dimensional posterior distribution of the dwell time *τ* _off_ and *τ* _on_. The color bar is a normalized probability density. The meanings of lines are the same as those in **(B). (D)** Posterior distribution of the burst frequency (BF) and burst size (BS). **(E-H)** The results of another example with the parameters *k*_off_ = 5, *r*_off_ = 3, *k*_on_ = 1, *r*_on_ = 0.8, *r*_syn_ = 30, *r*_deg_ = 1.

These results indicate that the BayesGTM inference of the GTM breaks the limitation that the dwell times only follow exponential distributions, thus enabling the correction of dynamic burst kinetics information from static single-cell snapshot data and the potential to extend it to genome-wide studies.

### 4.2 Experimental data

Next, we apply the BayesGTM inference procedure to mouse embryonic fibroblasts for each allele (C57×CAST) (Larsson, et al., 2019) to estimate transcriptional bursting parameters on a genome-wide scale. This scRNA-seq data was widely used to investigate genome-wide transcriptional burst kinetics (Larsson, et al., 2019; Larsson, et al., 2021; Luo, et al., 2022), which was sequenced based on smart-seq2 technology (UMI counts were used) and contained 10727 genes and 224 individual cells on each allele. For quality control, we filter out genes expressed in less than 50 cells and cells expressed in less than 2,000 genes. In addition, we remove genes with mean expression levels below 2 to ensure high expression of the inferred genes. Finally, we combine the two allele expressions together to eventually form a single cell matrix consisting of 2162 genes and 413 cells.

Interestingly, we observe that genes with the same average expression level have a different combination of burst frequency and burst size, consistent with previous reports (Larsson, et al., 2019; Suter, et al., 2011). This result implies gene expression may be regulated by diverse burst kinetic mechanisms (Figure 5A). In addition, we perform genome-wide burst kinetics inference for the same data based on the CTM, using a maximum likelihood estimation method. We discover that the estimated burst frequency and burst size keep a high positive correlation between GTM and CTM (p-value < 2.2×10-^16^, Figure 5B and 5C). Notably, we observe that the transcriptional burst kinetics inferred with the CTM would overestimate burst frequency (Figure 5B) and underestimate burst size (Figure 5C) on the genome-wide scale, consistent with the results for single genes in Figures 2D and 2H. Moreover, BayesGTM with an alternative definition of the burst frequency (1/⟨*τ*_off_⟩) also presents the same results (Supplementary Figure S3). These results suggest that the GTM, as an extension of the CTM, is more accurate in predicting burst kinetics, and BayesGTM has the capability to perform genome-wide studies on scRNA-seq data.

**Figure 5.**
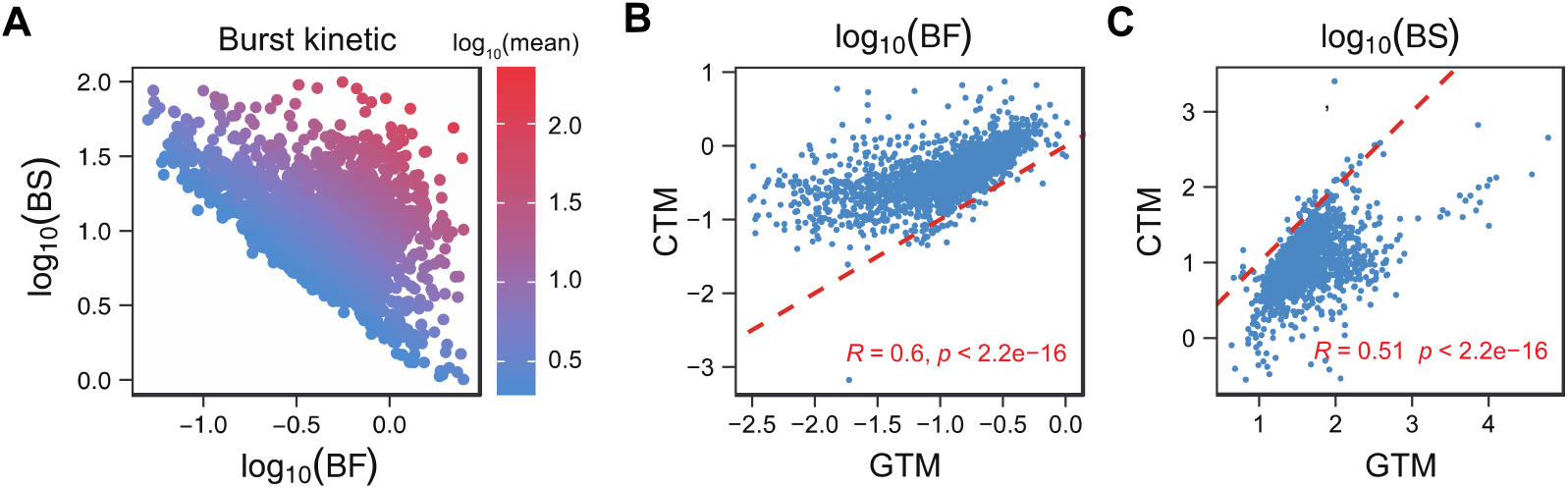
Genome-wide characteristics of transcriptional burst kinetics inferred from the scRNA-seq data of mouse embryonic fibroblasts. **(A)** Scatter plot of burst frequency (BF) and burst size (BS) inferred by BayesGTM, where the color bar represents the mean expression levels of each gene. **(B-C)** Scatter plots show the burst frequency **(B)** and burst size **(C)** estimated by GTM and CTM, which are correlated in the sense of Pearson correlation test (p-value < 2.2×10^−16^). The slope of red dashed lines equals 1.

## 5. Discussion

With the emergence of next-generation sequencing technologies, inferring gene expression burst kinetics on a genome-wide scale from static single-cell snapshot data is challenging in computational systems biology (Rodriguez and Larson, 2020). In previous studies (Kim and Marioni, 2013; Larsson, et al., 2019), gene expression models used for inference and analysis, e.g., CTM, have long relied on Markov assumptions. However, increasing experimental data show that the dwell time of states is not simply exponentially distributed (Dunham, et al., 2017; Harper, et al., 2011; Rodriguez, et al., 2019; Suter, et al., 2011), leading to the failure of the Markov approximation. There is thereby an urgent need but it is still a significant challenge to develop effective methods for modeling, analyzing, and inferring non-Markov gene expression models.

In this paper, we have developed the BayesGTM for inferring burst kinetics from scRNA-seq data based on GTM, which allows consideration of ON-OFF transitions with arbitrary dwell time distributions. We demonstrated that the CTM could not estimate the burst kinetics, although it can well fit the gene expression distributions sometimes. We theoretically derived the analytical solution for arbitrary order binomial moments of GTM, which in turn enables us to calculate the statistics of mRNA. We developed the BayesGTM inference procedure to infer transcriptional burst kinetics, using the summary statistic calculated by binomial moments. The results of the synthetic dataset show that our inference method can precisely estimate the burst frequency and burst size of the gene expression system as well as the average dwell time in ON and OFF states. Furthermore, we performed a genome-wide burst kinetics inference on the mouse embryonic fibroblasts scRNA-seq data with the BayesGTM. We found that the transcriptional burst kinetics inferred with the CTM would overestimate burst frequency and underestimate burst size.

The GTM and the corresponding BayesGTM are applicable to study burst kinetics on a genome-wide scale. First, the GTM is interpretable. The GTM, as an extension of the CTM, is a mechanistic model that considers the dwell times to be arbitrary distributions. And the model parameters, such as dwell times of OFF and ON states, transcription rates, and degradation rates, are measurable from experiments (Lammers, et al., 2020; Suter, et al., 2011). Second, the GTM is solvable. The arbitrary order binomial moments of mRNA’s distribution are theoretically derived. The model extends previous results. For example, the CTM is a special case of the GTM if *f*_off_ (*t*) and *f*_on_ (*t*) set to exponential distributions. Third, the inference method for GTM is scalable. The approximation of the probability distributions of the stochastic model is often computationally prohibitive. With the theoretical results derived, we used a “likelihood-free” approach by computing only low-order moments instead of the whole probability distribution. The efficiency of the BayesGTM facilitates the extension of individual genes to genome-wide study. Finally, our inference method can learn gene expression information from static snapshot data without time-resolved data.

Our work opens up several avenues for further research. From a modeling perspective, the GTM simplifies biological processes in several aspects. The GTM only considers the generalized dwell time distribution for OFF and ON states but does not consider the nonexponential waiting time for transcriptional and degradation processes. Many experimental data suggest that transcription (such as mRNA elongation, pause, and release (Dobrzyński and Bruggeman, 2009; Engl, et al., 2020; Tantale, et al., 2021)) and degradation (such as mRNA senescence (Decker and Parker, 1993; Keene, et al., 2001; Pedraza and Paulsson, 2008)) are also multi-step processes in cells. From a statistical inferring perspective, using GTM to interpret the experimental biological data has yet to be fully developed. First, the binomial moments obtained from our analysis are in the sense of a steady distribution (*t* ⟶∞) and not as a function of time *t*. Solving for the model’s temporal solutions facilitates applying the GTM to time-resolved data. Second, the inference based on the steady-state distribution still suffers from the unidentifiability of the parameters, which may depend on the properties that different dwell time distributions lead to the same static mRNA distribution.

Finally, we note that our approach is not limited to scRNA-seq data, but could also be useful for other kinds of single-cell data in which the probability distributions of mRNA can be estimated, for example, smFISH data. We expect the GTM and BayesGTM to facilitate an understanding of gene expression mechanisms from the enormous amount of biological data.

## Data availability

All the analysis results and inference code that support the findings of this study are available through GitHub (https://github.com/cellfate/BayesGTM).

## Reference

Alfa, A.S., and Rao, T.S. (2000) Supplementary variable technique in stochastic models. Probab. Eng. Inform. Sc., 14(2), 203–218.

Bertrand, E. et al. (1998) Localization of ASH1 mRNA particles in living yeast. Mol. Cell, 2(4), 437–445.

Blake, W.J. et al. (2006) Phenotypic consequences of promoter-mediated transcriptional noise. Mol. Cell, 24(6), 853–865.

Chao, J.A. et al. (2008) Structural basis for the coevolution of a viral RNA-protein complex. Nat. Struct. Mol. Biol., 15(1), 103–105.

Chen, S.Y. et al. (2020) Optogenetic control reveals differential promoter interpretation of transcription factor nuclear translocation dynamics. Cell Syst., 11(4), 336-353. e324.

Cox, D.R. The analysis of non-Markovian stochastic processes by the inclusion of supplementary variables. Cambridge, England: Cambridge University Press; 1955.

Daigle Jr, B.J. et al. (2015) Inferring single-cell gene expression mechanisms using stochastic simulation. Bioinformatics, 31(9), 1428–1435.

Decker, C.J., and Parker, R. (1993) A turnover pathway for both stable and unstable mRNAs in yeast: evidence for a requirement for deadenylation. Gene Dev., 7(8), 1632–1643.

Desai, R.V. et al. (2021) A DNA repair pathway can regulate transcriptional noise to promote cell fate transitions. Science, 373(6557), eabc6506.

Dobrzynski, M., and Bruggeman, F.J. (2009) Elongation dynamics shape bursty transcription and translation. Proc. Natl. Acad. Sci. U. S. A., 106(8), 2583–2588.

Dunham, L.S. et al. (2017) Asymmetry between activation and deactivation during a transcriptional pulse. Cell Syst., 5(6), 646–653.

Eldar, A., and Elowitz, M.B. (2010) Functional roles for noise in genetic circuits. Nature, 467(7312), 167–173.

Ellison, A.M. (1987) Effect of seed dimorphism on the density-dependent dynamics of experimental populations of Atriplex triangularis (Chenopodiaceae). Am. J. Bot., 74(8), 1280–1288.

Engl, C. et al. (2020) The route to transcription initiation determines the mode of transcriptional bursting in E. coli. Nat. Commun., 11(1), 2422.

Femino, A.M. et al. (1998) Visualization of single RNA transcripts in situ. Science, 280(5363), 585–590.

Franks, J.J. Handbook of Approximate Bayesian Computation. Boca Raton, Florida: CRC Press; 2020.

Fritzsch, C. et al. (2018) Estrogen-dependent control and cell-to-cell variability of transcriptional bursting. Mol. Syst. Biol., 14(2), e7678.

Fröhlich, F. et al. (2016) Inference for stochastic chemical kinetics using moment equations and system size expansion. PLoS Comput. Biol., 12(7), e1005030.

Fuda, N.J., Ardehali, M.B., and Lis, J.T. (2009) Defining mechanisms that regulate RNA polymerase II transcription in vivo. Nature, 461(7261), 186–192.

Harper, C.V. et al. (2011) Dynamic analysis of stochastic transcription cycles. PLoS Biol., 9(4), e1000607.

Jia, T., and Kulkarni, R.V. (2011) Intrinsic noise in stochastic models of gene expression with molecular memory and bursting. Phys. Rev. Lett., 106(5), 058102.

Jiang, Y., Zhang, N.R., and Li, M. (2017) SCALE: modeling allele-specific gene expression by single-cell RNA sequencing. Genome Biol., 18(1), 74.

Keene, J. et al. (2001) As examples accumulate of both ARE-bearing stable mRNAs and labile mRNAs lacking AREs, the ARE dogma has incre-mentally given way to alternative bona. Nat. Rev. Mol. Cell Biol, 2, 237–246.

Kim, J.K., and Marioni, J.C. (2013) Inferring the kinetics of stochastic gene expression from single-cell RNA-sequencing data. Genome Biol., 14(1), R7.

Klindziuk, A., and Kolomeisky, A.B. (2018) Theoretical investigation of transcriptional bursting: a multistate approach. J. Phys. Chem. B, 122(50), 11969–11977.

Kumar, N., Singh, A., and Kulkarni, R.V. (2015) Transcriptional bursting in gene expression: analytical results for general stochastic models. PLoS Comput. Biol., 11(10), e1004292.

Lammers, N.C. et al. (2020) A matter of time: Using dynamics and theory to uncover mechanisms of transcriptional bursting. Curr. Opin. Cell Biol., 67, 147–157.

Larson, D.R. et al. (2011) Real-time observation of transcription initiation and elongation on an endogenous yeast gene. Science, 332(6028), 475–478.

Larsson, A.J. et al. (2019) X-chromosome upregulation is driven by increased burst frequency. Nat. Struct. Mol. Biol., 26(10), 963–969.

Larsson, A.J. et al. (2019) Genomic encoding of transcriptional burst kinetics. Nature, 565(7738), 251–254.

Larsson, A.J. et al. (2021) Transcriptional bursts explain autosomal random monoallelic expression and affect allelic imbalance. PLoS Comput. Biol., 17(3), e1008772.

Lenive, O., Pd, W.K., and MP, H.S. (2016) Inferring extrinsic noise from single-cell gene expression data using approximate Bayesian computation. BMC Syst. Biol., 10(1), 81.

Luo, S. et al. (2022) Genome-wide inference reveals that feedback regulations constrain promoter-dependent transcriptional burst kinetics. bioRxiv https://doi.org/10.1101/2022.04.08.487618.

Ochab-Marcinek, A., and Tabaka, M. (2010) Bimodal gene expression in noncooperative regulatory systems. Proc. Natl. Acad. Sci. U. S. A., 107(51), 22096–22101.

Ochiai, H. et al. (2020) Genome-wide kinetic properties of transcriptional bursting in mouse embryonic stem cells. Sci. Adv., 6(25), eaaz6699.

Peccoud, J., and Ycart, B. (1995) Markovian modeling of gene-product synthesis. Theor. Popul. Biol., 48(2), 222–234.

Pedraza, J.M., and Paulsson, J. (2008) Effects of molecular memory and bursting on fluctuations in gene expression. Science, 319(5861), 339–343.

Picelli, S. et al. (2013) Smart-seq2 for sensitive full-length transcriptome profiling in single cells. Nat. Methods, 10(11), 1096–1098.

Raj, A. et al. (2006) Stochastic mRNA synthesis in mammalian cells. PLoS Biol., 4(10), e309.

Raj, A. et al. (2008) Imaging individual mRNA molecules using multiple singly labeled probes. Nat. Methods, 5(10), 877–879.

Rodriguez, J., and Larson, D.R. (2020) Transcription in living cells: molecular mechanisms of bursting. Annu. Rev. Biochem., 89, 189–212.

Rodriguez, J. et al. (2019) Intrinsic dynamics of a human gene reveal the basis of expression heterogeneity. Cell, 176(1-2), 213–226.

Sarkar, A., and Stephens, M. (2021) Separating measurement and expression models clarifies confusion in single-cell RNA sequencing analysis. Nat. Genet., 53(6), 770–777.

Schwabe, A., Rybakova, K.N., and Bruggeman, F.J. (2012) Transcription stochasticity of complex gene regulation models. Biophys. J., 103(6), 1152–1161.

Sepúlveda, L.A. et al. (2016) Measurement of gene regulation in individual cells reveals rapid switching between promoter states. Science, 351(6278), 1218–1222.

Shi, C., Jiang, Y., and Zhou, T. (2020) Queuing models of gene expression: Analytical distributions and beyond. Biophys. J., 119(8), 1606–1616.

Sisson, S.A., Fan, Y., and Tanaka, M.M. (2007) Sequential Monte Carlo without likelihoods. Proc. Natl. Acad. Sci. U. S. A., 104(6), 1760–1765.

Stinchcombe, A.R., Peskin, C.S., and Tranchina, D. (2012) Population density approach for discrete mRNA distributions in generalized switching models for stochastic gene expression. Phys. Rev. E, 85(6), 061919.

Stumpf, P.S. et al. (2017) Stem cell differentiation as a non-Markov stochastic process. Cell Syst., 5(3), 268–282.

Suter, D.M. et al. (2011) Mammalian genes are transcribed with widely different bursting kinetics. Science, 332(6028), 472–474.

Tantale, K. et al. (2021) Stochastic pausing at latent HIV-1 promoters generates transcriptional bursting. Nat. Commun., 12(1), 4503.

Tunnacliffe, E., and Chubb, J.R. (2020) What is a transcriptional burst? Trends. Genet., 36(4), 288–297.

Van Kampen, N.G. Stochastic processes in physics and chemistry. Elsevier; 1992.

Voss, T.C., and Hager, G.L. (2014) Dynamic regulation of transcriptional states by chromatin and transcription factors. Nat. Rev. Genet., 15(2), 69–81.

Vu, T.N. et al. (2016) Beta-Poisson model for single-cell RNA-seq data analyses. Bioinformatics, 32(14), 2128–2135.

Yu, J. et al. (2006) Probing gene expression in live cells, one protein molecule at a time. Science, 311(5767), 1600–1603.

Zechner, C. et al. (2012) Moment-based inference predicts bimodality in transient gene expression. Proc. Natl. Acad. Sci. U. S. A., 109(21), 8340–8345.

Zhang, J., Chen, L., and Zhou, T. (2012) Analytical distribution and tunability of noise in a model of promoter progress. Biophys. J., 102(6), 1247–1257.

Zhang, J., Nie, Q., and Zhou, T. (2016) A moment-convergence method for stochastic analysis of biochemical reaction networks. J. Chem. Phys., 144(19), 194109.

Zhang, J., and Zhou, T. (2014) Promoter-mediated transcriptional dynamics. Biophys. J., 106(2), 479–488.

Zhang, J., and Zhou, T. (2019) Markovian approaches to modeling intracellular reaction processes with molecular memory. Proc. Natl. Acad. Sci. U. S. A., 116(47), 23542–23550.

Zhang, J., and Zhou, T. (2019) Stationary moments, distribution conjugation and phenotypic regions in stochastic gene transcription. Math. Biosci. Eng., 16(5), 6134–6166.

Zheng, G.X. et al. (2017) Massively parallel digital transcriptional profiling of single cells. Nat. Commun., 8(1), 1–12.

Zhou, T., and Zhang, J. (2012) Analytical results for a multistate gene model. SIAM J. Appl. Math., 72(3), 789–818.

Zoller, B. et al. (2015) Structure of silent transcription intervals and noise characteristics of mammalian genes. Mol. Syst. Biol., 11(7), 823.

