## Supplementary Material for "Inferring transcriptional bursting kinetics from single-cell snapshot data using a generalized telegraph model"

### Supplementary Notes

#### 1. Analytical derivation of burst frequency and burst size (Eq. (2) and (3) in the main text)

##### *1.1 Burst frequency*

Burst frequency is defined as the reciprocal of the mean cycle time, where the cycle time is defined as the summation of OFF state and ON state dwell time per burst, i.e. $\tau_{\mathrm{off}}+\tau_{\mathrm{on}}$. The distribution of cycle time, denoted by $f_{\mathrm{cycle}}(t)$, is given by

$$f_{\mathrm{cycle}}(t)=\left(f_{\mathrm{off}}*f_{\mathrm{on}}\right)(t)=\int_0^t f_{\mathrm{off}}(t-t')f_{\mathrm{on}}(t')dt', \tag{S1}$$

where $*$ represents the convolution operator.

It is easy to obtain the burst frequency for GTM

$$\mathrm{BF}=\frac{1}{\langle\tau_{\mathrm{off}}+\tau_{\mathrm{on}}\rangle}=\frac{1}{\langle\tau_{\mathrm{off}}\rangle+\langle\tau_{\mathrm{on}}\rangle}, \tag{S2}$$

where $\langle\tau_{\mathrm{off}}\rangle=\int_0^\infty tf_{\mathrm{off}}(t)dt$ and $\langle\tau_{\mathrm{on}}\rangle=\int_0^\infty tf_{\mathrm{on}}(t)dt$ are the mean OFF and ON state dwell time, respectively. When $\langle\tau_{\mathrm{on}}\rangle\ll\langle\tau_{\mathrm{off}}\rangle$, Eq. (S2) reduces to the mean activation rate $r_{\mathrm{on}}=\langle\tau_{\mathrm{off}}\rangle^{-1}$, which is the classical definition of burst frequency (Larsson, et al., 2019; Peccoud and Ycart, 1995).

##### *1.2 Burst size*

Burst size is defined as the average number of mRNA molecules produced per burst. Let $f_{\mathrm{syn}}(t)$ be the distribution of waiting time between two successive transcription events. The probability that there are $n$ transcription events conditioned on a fixed duration time $t$ of the ON state is calculated as follow,

$$\begin{aligned}
&P\left(B=0\middle|\tau_{\mathrm{ON}}=t\right)=S_{\mathrm{syn}}(t),\\
&P\left(B=n\middle|\tau_{\mathrm{ON}}=t\right)=\left(f_{\mathrm{syn}}^{(n-1)}*S_{\mathrm{syn}}\right)(t)\\
&\qquad\qquad\qquad\quad=\int_0^t f_{\mathrm{syn}}^{(n-1)}(t')S_{\mathrm{syn}}(t-t')dt',\ n\ge 1,
\end{aligned} \tag{S3}$$

where $S_{\mathrm{syn}}(t)=\int_t^\infty f_{\mathrm{syn}}(t')dt'$ is the survival function for the waiting time between transcription events. For $0=t_0<t_1<\cdots<t_n=t'\le t$ with $n\ge 1$,

$$f_{\mathrm{syn}}^{(n-1)}(t')=\underbrace{\left(f_{\mathrm{syn}}*\cdots*f_{\mathrm{syn}}\right)}_{n-1}(t')=\int_0^{t'}\cdots\int_0^{t_2}\prod_{i=1}^n f_{\mathrm{syn}}(t_i-t_{i-1})dt_1\cdots dt_{n-1}, \tag{S4}$$

is the ($n-1$)-th convolution of $f_{\mathrm{syn}}(t')$. Here $f_{\mathrm{syn}}^{(0)}(t')=f_{\mathrm{syn}}(t')$. Eqs. (S3) with (S4) have a clear interpretation: the $i$-th transcription event occurs at time $t_i$ ($i=0,1,\cdots,n$), and there is no transcription in the remaining time $t-t_n$.

It is easy to prove $\tilde{S}(s)=\int_0^\infty\int_t^\infty e^{-st}f(t')dt'dt=\left(1-\tilde{f}(s)\right)/s$. Therefore, Eq. (S3) can be converted into

$$P\left(B=n|\tau_{\text{on}}=t\right)=\mathcal{L}^{-1}\left(\tilde{f}_{\text{syn}}^{n}(s)\frac{1-\tilde{f}_{\text{syn}}(s)}{s}\right),\ n\geq 0, \tag{S5}$$

where $\mathcal{L}^{-1}$ represents the inverse Laplace transform.

For exponential transcription process with rate $r_{\text{syn}}$, i.e. $f_{\text{syn}}(t)=r_{\text{syn}}e^{-r_{\text{syn}}t}$, the conditioned probability described by Eq. (S5) is a time-dependent Poisson distribution,

$$P\left(B=n|\tau_{\text{on}}=t\right)=\mathcal{L}^{-1}\left(\frac{r_{\text{syn}}^{n}}{\left(s+r_{\text{syn}}\right)^{n+1}}\right)=\frac{\left(r_{\text{syn}}t\right)^{n}}{n!}e^{-r_{\text{syn}}t},\ n\geq 0. \tag{S6}$$

Note that Eq. (S6) is the transient probability of the pure birth process. After integrating overall ON state dwell time $\tau_{\text{on}}$, we obtain

$$\begin{aligned}P(B=n)&=\int_0^\infty P\left(B=n|\tau_{\text{on}}=t\right)f_{\text{on}}(t)dt\\&=\int_0^\infty\frac{\left(r_{\text{syn}}t\right)^{n}}{n!}e^{-r_{\text{syn}}t}f_{\text{on}}(t)dt=\frac{r_{\text{syn}}^{n}}{n!}(-1)^{n}\frac{d^{n}\tilde{f}_{\text{on}}(s)}{ds^{n}}\bigg|_{s=r_{\text{syn}}}.\end{aligned} \tag{S7}$$

Then, the burst size of mRNA for GTM is given by

$$\begin{aligned}\text{BS}&=\sum_{n=0}^{\infty}bP(B=n)\\&=\sum_{n=0}^{\infty}\int_0^\infty n\frac{\left(r_{\text{syn}}t\right)^{n}}{n!}e^{-r_{\text{syn}}t}f_{\text{on}}(t)dt\\&=r_{\text{syn}}\int_0^\infty tf_{\text{on}}(t)dt=r_{\text{syn}}\langle\tau_{\text{on}}\rangle.\end{aligned} \tag{S8}$$

**2. Analytical derivation of binomial moments (Eq. (7) and (8) in the main text)**

To solve Eq. (6) with its boundary conditions, we define the probability generating functions for OFF state and ON state: $G_{\text{off}}(s+1,\tau)=\sum_{n=0}^{\infty}(s+1)^{n}p_{\text{off}}(n,\tau)$ and $G_{\text{on}}(s+1,\tau)=\sum_{n=0}^{\infty}(s+1)^{n}p_{\text{on}}(n,\tau)$, which both depend on elapsed time $\tau$. Then, Eq. (7) converts to the following partial differential equation for probability generating functions,

$$\begin{aligned}\frac{\partial G_{\text{off}}(s+1,\tau)}{\partial\tau}&=-H_{\text{off}}(\tau)G_{\text{off}}(s+1,\tau)-r_{\text{deg}}s\frac{\partial G_{\text{off}}(s+1,\tau)}{\partial s},\\\frac{\partial G_{\text{on}}(s+1,\tau)}{\partial\tau}&=-H_{\text{on}}(\tau)G_{\text{on}}(s+1,\tau)+r_{\text{syn}}sG_{\text{on}}(s+1,\tau)-r_{\text{deg}}s\frac{\partial G_{\text{on}}(s+1,\tau)}{\partial s},\end{aligned} \tag{S9}$$

with the corresponding boundary conditions

$$G_{\text{off}}(s+1,0)=\int_0^\infty G_{\text{on}}(s+1,\tau)H_{\text{on}}(\tau)d\tau,\\ G_{\text{on}}(s+1,0)=\int_0^\infty G_{\text{off}}(s+1,\tau)H_{\text{off}}(\tau)d\tau. \tag{S10}$$

A moment-convergence approach (Zhang, et al., 2016) is applied to calculate the mRNA stationary distribution and its raw or central moments here. First, we define elapsed time-dependent binomial moments for OFF state and ON state as

$$b_{n,\text{off}}(\tau)=\sum_{m\ge n}^{\infty}\binom{m}{n}p_{\text{off}}(m,\tau),\\ b_{n,\text{on}}(\tau)=\sum_{m\ge n}^{\infty}\binom{m}{n}p_{\text{on}}(m,\tau), \tag{S11}$$

respectively, where the symbol $\binom{m}{n}$ represents the combinatorial number. There are relationships between binomial moments and probability generating functions: $G_{\text{off}}(s+1,\tau)=\sum_{n=0}^{\infty}b_{n,\text{off}}(\tau)s^n$ and $G_{\text{on}}(s+1,\tau)=\sum_{n=0}^{\infty}b_{n,\text{on}}(\tau)s^n$. Thus, we can obtain the following ordinary differential equation for binomial moments,

$$\frac{db_{n,\text{off}}(\tau)}{d\tau}=-\left(nr_{\text{deg}}+H_{\text{off}}(\tau)\right)b_{n,\text{off}}(\tau),\\ \frac{db_{n,\text{on}}(\tau)}{d\tau}=-\left(nr_{\text{deg}}+H_{\text{on}}(\tau)\right)b_{n,\text{on}}(\tau)+r_{\text{syn}}b_{n-1,\text{on}}(\tau). \tag{S12}$$

Here we define $b_{-1,\text{on}}(\tau)\equiv 0$. The boundary conditions for binomial moments are given by

$$b_{n,\text{off}}(0)=\int_0^\infty b_{n,\text{on}}(\tau)H_{\text{on}}(\tau)d\tau,\\ b_{n,\text{on}}(0)=\int_0^\infty b_{n,\text{off}}(\tau)H_{\text{off}}(\tau)d\tau. \tag{S13}$$

Solving Eq. (S12) yields

$$b_{n,\text{off}}(\tau)=S_{\text{off}}(\tau)e^{-nr_{\text{deg}}\tau}b_{n,\text{off}}(0),\\ b_{n,\text{on}}(\tau)=S_{\text{on}}(\tau)e^{-nr_{\text{deg}}\tau}b_{n,\text{on}}(0)+r_{\text{syn}}e^{-nr_{\text{deg}}\tau}A_n(\tau)S_{\text{on}}(\tau), \tag{S14}$$

where $A_n(\tau)=\int_0^\tau e^{nr_{\text{deg}}\tau'}b_{n-1,\text{on}}(\tau')S_{\text{on}}^{-1}(\tau')d\tau'$.

We denote $f_n=\int_0^\infty e^{-nr_{\text{deg}}\tau}A_n(\tau)f_{\text{on}}(\tau)d\tau$ and $S_n=\int_0^\infty e^{-nr_{\text{deg}}\tau}A_n(\tau)S_{\text{on}}(\tau)d\tau$ for the simplicity of calculation. Note that $A_0(\tau)=f_0=S_0=0$. Substituting Eq. (S14) into Eq. (S13), we have

$$b_{n,\text{off}}(0)=\tilde{f}_{\text{on}}(n\delta)b_{n,\text{on}}(0)+r_{\text{syn}}f_n,\\ b_{n,\text{on}}(0)=\tilde{f}_{\text{off}}(n\delta)b_{n,\text{off}}(0). \tag{S15}$$

Then, the boundary values can be solved as

$$b_{0,\text{off}}(0)=b_{0,\text{on}}(0), \tag{S16}$$

and

$$b_{n,\text{off}}(0)=\frac{r_{\text{syn}}f_n}{1-\tilde f_{\text{off}}(nr_{\text{deg}})\tilde f_{\text{on}}(nr_{\text{deg}})},\ b_{n,\text{on}}(0)=\frac{\tilde f_{\text{off}}(nr_{\text{deg}})r_{\text{syn}}f_n}{1-\tilde f_{\text{off}}(nr_{\text{deg}})\tilde f_{\text{on}}(nr_{\text{deg}})},\ n=1,2,\cdots. \tag{S17}$$

Note that $\tilde f_{\text{off}}(0)=\int_0^\infty f_{\text{off}}(t)dt=1$ and $\tilde f_{\text{on}}(0)=\int_0^\infty f_{\text{on}}(t)dt=1$.

The $n$-th binomial moment of the mRNA stationary distribution is calculated according to Eq. (S14),

$$\begin{aligned}b_n&=\int_0^\infty b_{n,\text{off}}(\tau)d\tau+\int_0^\infty b_{n,\text{on}}(\tau)d\tau\\&=b_{n,\text{off}}(0)\tilde S_{\text{off}}(nr_{\text{deg}})+b_{n,\text{on}}(0)\tilde S_{\text{on}}(nr_{\text{deg}})+r_{\text{syn}}S_n\end{aligned} \tag{S18}$$

for $n=0,1,2,\cdots$. Since $b_0=\sum_{m=0}^{\infty}\int_0^\infty\left(p_{\text{off}}(m,\tau)+p_{\text{on}}(m,\tau)\right)d\tau=1$, it is easy to know

$$b_{0,\text{off}}(0)=b_{0,\text{on}}(0)=\left(\tilde S_{\text{off}}(0)+\tilde S_{\text{on}}(0)\right)^{-1}=\text{BF}. \tag{S19}$$

Note that $\tilde S_{\text{off}}(0)=\int_0^\infty S_{\text{off}}(t)dt=\langle\tau_{\text{off}}\rangle$ and $\tilde S_{\text{on}}(0)=\int_0^\infty S_{\text{on}}(t)dt=\langle\tau_{\text{on}}\rangle$.

Eq. (S18) remains unknown expressions of $f_n$ and $S_n$, so we need to calculate $A_n(\tau)$ first. According to Eq. (S14),

$$A_n(\tau)=\frac{b_{n-1,\text{on}}(0)}{\delta}\left(e^{r_{\text{deg}}\tau}-1\right)+\mu\int_0^\tau e^{r_{\text{deg}}\tau'}A_{n-1}(\tau')d\tau', \tag{S20}$$

for $n=1,2,\cdots$. By the mathematical induction and the additional fact that

$$\int_0^\tau e^{r_{\text{deg}}\tau'}\left(e^{r_{\text{deg}}\tau'}-1\right)^n d\tau'=\frac{1}{(n+1)r_{\text{deg}}}\left(e^{r_{\text{deg}}\tau}-1\right)^{n+1},\ n=0,1,2,\cdots, \tag{S21}$$

we can prove

$$A_n(\tau)=\frac{1}{r_{\text{syn}}}\sum_{i=0}^{n-1}\left(\frac{r_{\text{syn}}}{r_{\text{deg}}}\right)^{n-i}\frac{b_{i,\text{on}}(0)}{(n-i)!}\left(e^{r_{\text{deg}}\tau}-1\right)^{n-i}. \tag{S22}$$

Therefore, $f_n$ and $S_n$ are obtained by iterative formulae of $b_{i,\text{on}}(0)$,

$$\begin{aligned}f_n&=\frac{1}{r_{\text{syn}}}\sum_{i=0}^{n-1}\left(\frac{r_{\text{syn}}}{r_{\text{deg}}}\right)^{n-i}\frac{b_{i,\text{on}}(0)}{(n-i)!}\int_0^\infty e^{-nr_{\text{deg}}\tau}\left(e^{r_{\text{deg}}\tau}-1\right)^{n-i}f_{\text{on}}(\tau)d\tau,\\S_n&=\frac{1}{r_{\text{syn}}}\sum_{i=0}^{n-1}\left(\frac{r_{\text{syn}}}{r_{\text{deg}}}\right)^{n-i}\frac{b_{i,\text{on}}(0)}{(n-i)!}\int_0^\infty e^{-nr_{\text{deg}}\tau}\left(e^{r_{\text{deg}}\tau}-1\right)^{n-i}S_{\text{on}}(\tau)d\tau.\end{aligned} \tag{S23}$$

In addition,

$$\int_0^\infty e^{-nr_{\deg}\tau}\left(e^{r_{\deg}\tau}-1\right)^{n-i} f_{\text{on}}(\tau)d\tau = -nr_{\deg}\int_0^\infty e^{-nr_{\deg}\tau}\left(e^{r_{\deg}\tau}-1\right)^{n-i} S_{\text{on}}(\tau)d\tau \\ +(n-i)r_{\deg}\int_0^\infty e^{-(n-1)r_{\deg}\tau}\left(e^{r_{\deg}\tau}-1\right)^{n-1-i} S_{\text{on}}(\tau)d\tau. \tag{S24}$$

Therefore, $f_n$ and $S_n$ have the following relationship,

$$f_n = -nr_{\deg}S_n + \sum_{i=0}^{n-1}\left(\frac{r_{\text{syn}}}{r_{\deg}}\right)^{n-1-i}\frac{b_{i,\text{on}}(0)}{(n-1-i)!}\int_0^\infty e^{-(n-1)r_{\deg}\tau}\left(e^{r_{\deg}\tau}-1\right)^{n-1-i} S_{\text{on}}(\tau)d\tau \\ = -nr_{\deg}S_n + \sum_{i=0}^{n-1}\left(\frac{r_{\text{syn}}}{r_{\deg}}\right)^{n-1-i}\frac{b_{i,\text{on}}(0)}{(n-1-i)!}\sum_{j=0}^{n-1-i}\binom{n-1-i}{j}(-1)^{n-1-i-j}\tilde{S}_{\text{on}}\left((n-1-j)r_{\deg}\right). \tag{S25}$$

Using this relationship, the $n$-th ($n=1,2,\cdots$) binomial moment $b_n$ has an iterative formula of $b_{i,\text{on}}(0)$,

$$b_n = \frac{r_{\text{syn}}}{r_{\deg}}\frac{f_n}{n} + r_{\text{syn}}S_n = \frac{1}{n}\sum_{i=0}^{n-1}\left(\frac{r_{\text{syn}}}{r_{\deg}}\right)^{n-i}\frac{b_{i,\text{on}}(0)}{(n-1-i)!}\sum_{j=0}^{n-1-i}\binom{n-1-i}{j}(-1)^{n-1-i-j}\tilde{S}_{\text{on}}\left((n-1-j)r_{\deg}\right). \tag{S26}$$

Note that a recursive form of $b_{n,\text{on}}(0)$ for $n=1,2,\cdots$ can be obtained by substituting the first equation of Eq. (S23) into the second equation of Eq. (S17)

$$b_{n,\text{on}}(0) = \frac{\tilde{f}_{\text{off}}(nr_{\deg})}{1-\tilde{f}_{\text{off}}(nr_{\deg})\tilde{f}_{\text{on}}(nr_{\deg})}\sum_{i=0}^{n-1}\left(\frac{r_{\text{syn}}}{r_{\deg}}\right)^{n-i}\frac{b_{i,\text{on}}(0)}{(n-i)!}\int_0^\infty e^{-nr_{\deg}\tau}\left(e^{r_{\deg}\tau}-1\right)^{n-i} f_{\text{on}}(\tau)d\tau \\ = \frac{\tilde{f}_{\text{off}}(nr_{\deg})}{1-\tilde{f}_{\text{off}}(nr_{\deg})\tilde{f}_{\text{on}}(nr_{\deg})}\sum_{i=0}^{n-1}\left(\frac{r_{\text{syn}}}{r_{\deg}}\right)^{n-i}\frac{b_{i,\text{on}}(0)}{(n-i)!}\sum_{j=0}^{n-i}\binom{n-i}{j}(-1)^{n-i-j}\tilde{f}_{\text{on}}\left((n-j)r_{\deg}\right). \tag{S27}$$

In order to extract the features of Poisson distribution, we define $C_n = \left(r_{\text{syn}}/r_{\deg}\right)^{-n} n! b_{n,\text{on}}(0)$. Then,

$$C_n = \frac{\tilde{f}_{\text{off}}(nr_{\deg})}{1-\tilde{f}_{\text{off}}(nr_{\deg})\tilde{f}_{\text{on}}(nr_{\deg})}\sum_{i=0}^{n-1}\binom{n}{i}C_i\sum_{j=0}^{n-i}\binom{n-i}{j}(-1)^{n-i-j}\tilde{f}_{\text{on}}\left((n-j)r_{\deg}\right). \tag{S28}$$

for $n=1,2,\cdots$, and $C_0 = \text{BF}$. Eq. (S26) converts to

$$b_n = \frac{1}{n!}\left(\frac{r_{\text{syn}}}{r_{\deg}}\right)^n\sum_{i=0}^{n-1}\binom{n-1}{i}C_i\sum_{j=0}^{n-1-i}\binom{n-1-i}{j}(-1)^{n-1-i-j}\tilde{S}_{\text{on}}\left((n-1-j)r_{\deg}\right). \tag{S29}$$

The mRNA stationary distribution can be reconstructed with a reconstruction formula (Zhang, et al., 2016),

$$P(n) = \sum_{m=n}^{\infty}(-1)^{m-n}\binom{m}{n}b_m. \tag{S30}$$

#### 3. Simplification of Eq. (7) for some special cases

##### *3.1 Arbitrary OFF state dwell time and exponential ON state dwell time*

In particular, assume that ON state dwell time is exponential, i.e. $f_{\text{on}}(t) = \langle\tau_{\text{on}}\rangle^{-1}S_{\text{on}}(t) = \langle\tau_{\text{on}}\rangle^{-1}e^{-t/\langle\tau_{\text{on}}\rangle}$. For any fixed $n \geq 1$, since $\tilde{f}_{\text{on}}(nr_{\deg}) = 1/\left(nr_{\deg}\langle\tau_{\text{on}}\rangle+1\right)$ and $\tilde{S}_{\text{on}}(nr_{\deg}) = \langle\tau_{\text{on}}\rangle\tilde{f}_{\text{on}}(nr_{\deg})$, Eq. (S29) becomes

$$b_n = \left(\frac{r_{\text{syn}}}{r_{\text{deg}}}\right)^n \frac{\langle \tau_{\text{on}} \rangle}{n!} \frac{C_{n-1}}{\tilde{f}_{\text{off}}\left(\left(n-1\right) r_{\text{deg}}\right)}, \tag{S31}$$

and

$$\begin{aligned} C_n &= \frac{\tilde{f}_{\text{off}}\left(nr_{\text{deg}}\right)\left(nr_{\text{deg}}\langle \tau_{\text{on}} \rangle + 1\right)}{nr_{\text{deg}}\langle \tau_{\text{on}} \rangle + 1 - \tilde{f}_{\text{off}}\left(nr_{\text{deg}}\right)} \sum_{i=0}^{n-1} \binom{n}{i} C_i \sum_{j=0}^{n-i} \binom{n-i}{j} \frac{(-1)^{n-i-j}}{(n-j) r_{\text{deg}} \langle \tau_{\text{on}} \rangle + 1} \\ &= \frac{\tilde{f}_{\text{off}}\left(nr_{\text{deg}}\right) r_{\text{deg}} \langle \tau_{\text{on}} \rangle}{nr_{\text{deg}}\langle \tau_{\text{on}} \rangle + 1 - \tilde{f}_{\text{off}}\left(nr_{\text{deg}}\right)} \sum_{i=0}^{n-1} \binom{n}{i} C_i \sum_{j=1}^{n-i} \binom{n-i}{j} \frac{(-1)^{n-i-j} j}{(n-j) r_{\text{deg}} \langle \tau_{\text{on}} \rangle + 1}. \end{aligned} \tag{S32}$$

For any $i \in \{0, 1, \cdots, n-1\}$, we can prove

$$\sum_{j=1}^{n-i} \binom{n-i}{j} \frac{(-1)^{n-i-j} j}{(n-j) r_{\text{deg}} \langle \tau_{\text{on}} \rangle + 1} = \frac{(n-i)! \left(r_{\text{deg}} \langle \tau_{\text{on}} \rangle\right)^{n-i-1}}{\prod_{j=i}^{n-1} \left(jr_{\text{deg}} \langle \tau_{\text{on}} \rangle + 1\right)} \tag{S33}$$

by the mathematical induction. Thus, Eq. (S32) converts to

$$C_n = \frac{\tilde{f}_{\text{off}}\left(nr_{\text{deg}}\right)}{nr_{\text{deg}}\langle \tau_{\text{on}} \rangle + 1 - \tilde{f}_{\text{off}}\left(nr_{\text{deg}}\right)} \frac{n! \left(r_{\text{deg}} \langle \tau_{\text{on}} \rangle\right)^n}{\prod_{j=1}^{n-1} \left(jr_{\text{deg}} \langle \tau_{\text{on}} \rangle + 1\right)} \sum_{i=0}^{n-1} \frac{\prod_{j=1}^{i-1} \left(jr_{\text{deg}} \langle \tau_{\text{on}} \rangle + 1\right)}{i! \left(r_{\text{deg}} \langle \tau_{\text{on}} \rangle\right)^i} C_i. \tag{S34}$$

Specially, Eq. (S34) implies that

$$\begin{aligned} \sum_{i=0}^{n} \frac{\prod_{j=1}^{i-1} \left(jr_{\text{deg}} \langle \tau_{\text{on}} \rangle + 1\right)}{i! \left(r_{\text{deg}} \langle \tau_{\text{on}} \rangle\right)^i} C_i &= \frac{nr_{\text{deg}} \langle \tau_{\text{on}} \rangle + 1}{nr_{\text{deg}}\langle \tau_{\text{on}} \rangle + 1 - \tilde{f}_{\text{off}}\left(nr_{\text{deg}}\right)} \sum_{i=0}^{n-1} \frac{\prod_{j=1}^{i-1} \left(jr_{\text{deg}} \langle \tau_{\text{on}} \rangle + 1\right)}{i! \left(r_{\text{deg}} \langle \tau_{\text{on}} \rangle\right)^i} C_i \\ &= \cdots \\ &= C_0 \prod_{j=1}^{n} \frac{jr_{\text{deg}} \langle \tau_{\text{on}} \rangle + 1}{jr_{\text{deg}} \langle \tau_{\text{on}} \rangle + 1 - \tilde{f}_{\text{off}}\left(jr_{\text{deg}}\right)}. \end{aligned} \tag{S35}$$

Then, $C_n$ has a concise expression as follow,

$$C_n = \frac{n! \left(r_{\text{deg}} \langle \tau_{\text{on}} \rangle\right)^n \tilde{f}_{\text{off}}\left(nr_{\text{deg}}\right)}{\left(\langle \tau_{\text{off}} \rangle + \langle \tau_{\text{on}} \rangle\right) \prod_{i=1}^{n} \left[ir_{\text{deg}} \langle \tau_{\text{on}} \rangle + 1 - \tilde{f}_{\text{off}}\left(ir_{\text{deg}}\right)\right]} = \frac{\langle \tau_{\text{on}} \rangle^n \tilde{f}_{\text{off}}\left(nr_{\text{deg}}\right)}{\prod_{i=0}^{n-1} \left[\langle \tau_{\text{on}} \rangle + \tilde{S}_{\text{off}}\left(ir_{\text{deg}}\right)\right]}. \tag{S36}$$

Therefore, the $n$-th binomial moment $b_n$ is given by substituting Eq. (S36) into Eq. (S31),

$$b_n = \left(\frac{r_{\text{syn}}}{r_{\text{deg}}}\right)^n \frac{\langle \tau_{\text{on}} \rangle^n}{n!} \frac{1}{\prod_{i=0}^{n-1} \left[\langle \tau_{\text{on}} \rangle + \tilde{S}_{\text{off}}\left(ir_{\text{deg}}\right)\right]}. \tag{S37}$$

Eq. (S37) is the same as the result derived in (Shi, et al., 2020).

#### *3.2 Exponential OFF state and ON state dwell time (CTM)*

For CTM, both OFF state and ON state dwell time are exponential. Assume that $f_{\text{on}}(t) = \langle \tau_{\text{on}} \rangle^{-1} e^{-t/\langle \tau_{\text{on}} \rangle}$ and $f_{\text{off}}(t) = \langle \tau_{\text{off}} \rangle^{-1} e^{-t/\langle \tau_{\text{off}} \rangle}$. Then, Eq. (S37) becomes

$$b_n = \left(\frac{r_{\text{syn}}}{r_{\deg}}\right)^n \frac{\langle \tau_{\text{on}} \rangle^n}{n!} \prod_{i=0}^{n-1} \left[ \langle \tau_{\text{on}} \rangle + \frac{\langle \tau_{\text{off}} \rangle}{i r_{\deg} \langle \tau_{\text{off}} \rangle + 1} \right]^{-1}$$
$$= \frac{1}{n!} \left(\frac{r_{\text{syn}}}{r_{\deg}}\right)^n \frac{\Gamma\left(\frac{r_{\text{on}}}{r_{\deg}} + n\right)}{\Gamma\left(\frac{r_{\text{on}}}{r_{\deg}}\right)} \frac{\Gamma\left(\frac{r_{\text{on}}}{r_{\deg}} + \frac{r_{\text{off}}}{r_{\deg}}\right)}{\Gamma\left(\frac{r_{\text{on}}}{r_{\deg}} + \frac{r_{\text{off}}}{r_{\deg}} + n\right)}, \tag{S38}$$

where $\Gamma(x)$ is the Gamma function. Note that $\prod_{i=0}^{n-1}(i+x) = \Gamma(x+n)/\Gamma(x)$ for any nonnegative integer $n$. Here $r_{\text{on}} = \langle \tau_{\text{off}} \rangle^{-1}$ is the mean activation rate, and $r_{\text{off}} = \langle \tau_{\text{on}} \rangle^{-1}$ is the mean inactivation rate.

According to Eq. (S30), the mRNA stationary distribution for CTM is

$$P(n) = \sum_{m=n}^{\infty} (-1)^{m-n} \binom{m}{n} \frac{1}{m!} \left(\frac{r_{\text{syn}}}{r_{\deg}}\right)^m \frac{\Gamma\left(\frac{r_{\text{on}}}{r_{\deg}} + m\right)}{\Gamma\left(\frac{r_{\text{on}}}{r_{\deg}}\right)} \frac{\Gamma\left(\frac{r_{\text{on}}}{r_{\deg}} + \frac{r_{\text{off}}}{r_{\deg}}\right)}{\Gamma\left(\frac{r_{\text{on}}}{r_{\deg}} + \frac{r_{\text{off}}}{r_{\deg}} + m\right)}$$
$$= \frac{1}{n!} \frac{\Gamma\left(\frac{r_{\text{on}}}{r_{\deg}} + n\right)}{\Gamma\left(\frac{r_{\text{on}}}{r_{\deg}}\right)} \frac{\Gamma\left(\frac{r_{\text{on}}}{r_{\deg}} + \frac{r_{\text{off}}}{r_{\deg}}\right)}{\Gamma\left(\frac{r_{\text{on}}}{r_{\deg}} + \frac{r_{\text{off}}}{r_{\deg}} + n\right)} \left(\frac{r_{\text{syn}}}{r_{\deg}}\right)^n {}_1F_1\left(\frac{r_{\text{on}}}{r_{\deg}} + n, \frac{r_{\text{on}}}{r_{\deg}} + \frac{r_{\text{off}}}{r_{\deg}} + n, -\frac{r_{\text{syn}}}{r_{\deg}}\right). \tag{S39}$$

where ${}_1F_1(x,y,z) = \sum_{n=0}^{\infty} \Gamma(x+n)\Gamma(y) z^n / n!\Gamma(x)\Gamma(y+n)$ is the confluent hypergeometric function of the first kind. This distribution for CTM has been derived in (Peccoud and Ycart, 1995; Raj, et al., 2006) and is equivalent to a Poisson-Beta distribution (Vu, et al., 2016).

### Supplementary Tables

**Table S1**

**Algorithm 1: Simulation of GTM**

**Input:**

1. Number of mRNA molecules $M \geq 0$.
2. Gene state $G \in \{\mathrm{OFF}, \mathrm{ON}\}$.
3. ON state dwell time distribution $f_{\mathrm{on}}(t)$ and OFF state dwell time distribution $f_{\mathrm{off}}(t)$.
4. Synthesis rate $r_{\mathrm{syn}} > 0$ and degradation rate $r_{\mathrm{deg}} > 0$.

**Sampling:**

1. Initialize $t = 0$, $M = 0$, and $G = \mathrm{OFF}$;
2. For $t = 0$ to $T$ do repeat:
   - I. If $G = \mathrm{OFF}$,
     - a. Sample the dwell time $\tau_{\mathrm{off}}$ at OFF state from $f_{\mathrm{off}}(t)$;
     - b. Set $\tau_{\mathrm{off}}^{\mathrm{end}} = t + \tau_{\mathrm{off}}$;
     - c. While $t < \tau_{\mathrm{off}}^{\mathrm{end}}$ do repeat:
       - c1. Set the reaction propensity $a_1 = M r_{\mathrm{deg}}$ for degradation;
       - c2. Sample one random value $r_1$ from a uniform distribution $U(0,1)$;
       - c3. Set $\tau = -\ln r_1 / a_1$, and $t = t + \tau$;
       - c4. If $t < \tau_{\mathrm{off}}^{\mathrm{end}}$, set $M = M - 1$;
     - d. Set $t = \tau_{\mathrm{off}}^{\mathrm{end}}$, and $G = \mathrm{ON}$;
   - II. If $G = \mathrm{ON}$,
     - a. Sample the dwell time $\tau_{\mathrm{on}}$ at ON state from $f_{\mathrm{on}}(t)$;
     - b. Set $\tau_{\mathrm{on}}^{\mathrm{end}} = t + \tau_{\mathrm{on}}$;
     - c. While $t < \tau_{\mathrm{on}}^{\mathrm{end}}$ do repeat:
       - c1. Set reaction propensities $a_1 = r_{\mathrm{syn}}$ for synthesis, $a_2 = M r_{\mathrm{deg}}$ for degradation;
       - c2. Sample two random values $r_1$, $r_2$ from a uniform distribution $U(0,1)$;
       - c3. Set $\tau = -\ln r_1 / (a_1 + a_2)$;
       - c4. Choose $q \in \{1,2\}$ to satisfy $\sum_{i=1}^{q-1} a_i \leq r_2 (a_1 + a_2) < \sum_{i=1}^{q} a_i$, and $t = t + \tau$;
       - c5. If $t < \tau_{\mathrm{on}}^{\mathrm{end}}$, If $q = 1$, set $M = M + 1$; If $q = 2$, set $M = M - 1$;
     - d. Set $t = \tau_{\mathrm{on}}^{\mathrm{end}}$, and $G = \mathrm{OFF}$.

**Output:**

Obtain a desired trajectory $(M, G, t)$.

**Table S2**

| **Algorithm 2**: **BayesGTM** |
|---|
| **Input:** |
| 1. Observational data $\mathbf{y}_{\text{obs}}$ . |
| 2. A pre-defined GTM with parameters $\boldsymbol{\theta}$ including $f_{\text{off}}(t)$, $f_{\text{on}}(t)$, $r_{\text{syn}}$ and $r_{\text{deg}}$ . |
| 2. Prior distribution $\pi(\boldsymbol{\theta})$. |
| 3. Number of samples $N > 0$. |
| 4. A summary statistics $\mathbf{s}_{\text{obs}} = S(\mathbf{y}_{\text{obs}})$, and $\mathbf{s}_{\text{model}} = S(\boldsymbol{\theta})$ expressed by binomial moments Eq (7). |
| 5. A discrepancy metrics $\rho(\mathbf{s}_{\text{model}}, \mathbf{s}_{\text{obs}})$ . |
| 6. A threshold values $\varepsilon_0$ . |
| 7. Proposal distribution $K_t(\boldsymbol{\theta} \vert \boldsymbol{\theta}')$ . |
| **Sampling:** |
| 1. Initialize $t = 0$ and weights $w_t^{(i)} = 1/N$ for $i = 1, \ldots, N$; |
| 2. For $t = 0$ to $T$ do repeat: |
| a. Set $i = 0$ |
| b. If $t = 0$, sample $\boldsymbol{\theta}$ from prior $\pi(\boldsymbol{\theta})$; |
| If $t \neq 0$, sample $\boldsymbol{\theta}'$ from $\{\boldsymbol{\theta}_{t-1}^{(i)}, w_{t-1}^{(i)}\}$ and $\boldsymbol{\theta}$ from $K_t(\boldsymbol{\theta} \vert \boldsymbol{\theta}')$; |
| c. Compute summary statistics of GTM $\mathbf{s}_{\text{model}} = S(\boldsymbol{\theta})$; |
| d. If $t = 0$, compute $\varepsilon_0^{(i)} = \rho(\mathbf{s}_{\text{model}}, \mathbf{s}_{\text{obs}})$; |
| d1. If $\varepsilon_0^{(i)} \leq \varepsilon_0$ , $\boldsymbol{\theta}_0^{(i)} = \boldsymbol{\theta}$ ; |
| d2. If $\varepsilon_0^{(i)} > \varepsilon_0$ , back to step b. |
| If $t \neq 0$, compute $\varepsilon_t^{(i)} = \rho(\mathbf{s}_{\text{model}}, \mathbf{s}_{\text{obs}})$ and $\varepsilon_t = median\left(\left\{\varepsilon_{t-1}^{(1)}, \ldots, \varepsilon_{t-1}^{(N)}\right\}\right)$ |
| d3. If $\varepsilon_t^{(i)} \leq \varepsilon_t$ , $\boldsymbol{\theta}_t^{(i)} = \boldsymbol{\theta}$ and weights $w_t^{(i)} = \pi(\boldsymbol{\theta}_t^{(i)}) \Big/ \sum_{j=1}^{N} w_{t-1}^{(j)} K_t(\boldsymbol{\theta}_t^{(i)} \vert \boldsymbol{\theta}_{t-1}^{(i)})$ |
| d4. If $\varepsilon_t^{(i)} > \varepsilon_t$ , back to step b. |
| e. set $i = i + 1$; |
| f. If $i < N$, back to step b. |
| **Output:** |
| Obtain a set of parameter vectors $\{\boldsymbol{\theta}_T^{(1)}, \ldots, \boldsymbol{\theta}_T^{(N)}\}$ from the parameter posterior distribution. |

### Supplementary Figures

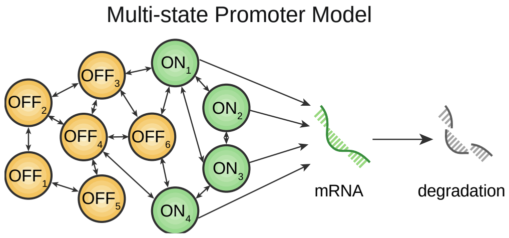

**Figure S1** The multi-state promoter model contains multiple OFF states ($OFF_1$,…,$OFF_6$, for example) and multiple ON states ($ON_1$,…,$ON_m$, for example), and states can switch arbitrarily with each other. The mRNA can be transcribed only in the ON state, and can be degraded in any state.

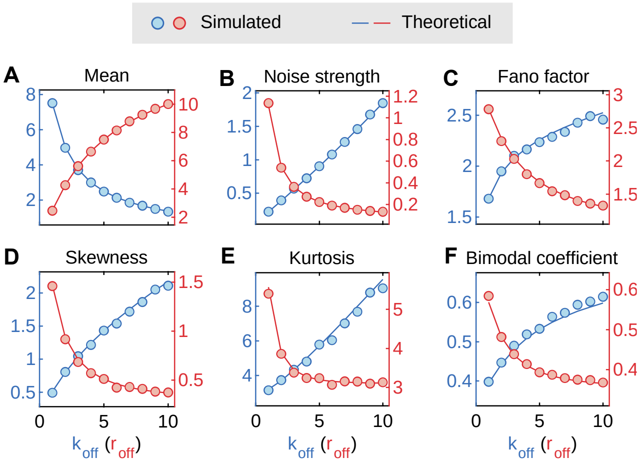

**Figure S2** Influence of the shape parameter $k_{\text{off}}$ (blue) and rate parameter $r_{\text{off}}$ (red) of the gamma dwell time distribution on the Mean (A), Noise strength (B), Fano factor (C), Skewness (D), Kurtosis (E) and Bimodal coefficient (F), where the solid lines correspond to theoretical values and the circles to the mean of 50 simulated values obtained by simulation algorithm of the GTM.

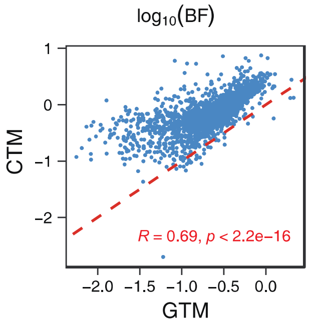

**Figure S3** Relationship (Scatter plot) between burst frequency (BF, defined by $1/\langle \tau_{\text{off}} \rangle$ ) of CTM and GTM, which are correlated in the sense of Pearson correlation test (p-value < $2.2\times10^{-16}$). The slope of red dashed lines equals 1.

### References

Larsson, A.J., *et al.* (2019) Genomic encoding of transcriptional burst kinetics. *Nature*, **565**(7738), 251-254.

Peccoud, J. and Ycart, B. (1995) Markovian modeling of gene-product synthesis. *Theor. Popul. Biol.*, **48**(2), 222-234.

Raj, A., *et al.* (2006) Stochastic mRNA synthesis in mammalian cells. *PLoS Biol.*, **4**(10), e309.

Shi, C., Jiang, Y. and Zhou, T. (2020) Queuing models of gene expression: Analytical distributions and beyond. *Biophys. J.*, **119**(8), 1606-1616.

Vu, T.N., *et al.* (2016) Beta-Poisson model for single-cell RNA-seq data analyses. *Bioinformatics*, **32**(14), 2128-2135.

Zhang, J., Nie, Q. and Zhou, T. (2016) A moment-convergence method for stochastic analysis of biochemical reaction networks. *J. Chem. Phys.*, **144**(19), 194109.
